# Unscheduled DNA synthesis reveals a DNA repair hotspot and biomarker of somatic instability at the expanded-CAG repeat tract in the huntingtin gene

**DOI:** 10.64898/2026.07.29.741355

**Authors:** Judith Praena Fernandez, Annette Gärtner, Jesús Perez-Perez, Sarah A. Cumming, Jia Feng, Katherine van Belois, Kangning He, Denny Yang, Carla Castells-Esteve, Maria Bergé-Gardeñes, Anna L. Bergander, Krzysztof P. Lubieniecki, Mihai Miclăuș, Nicolas Fissolo, Manuel Comabella, Alexander Vogt, Lukas Gebauer, Fatima Cavaleri, Christin Voelkner, Anastasia Illarionova, Hinal Limbani, Gina Marke, Richard Z. Chen, Brinda C. Prasad, Thomas F. Vogt, Dylan A. Reid, Ricardo Mouro Pinto, Petar Podlesniy, Jaime Kulisevsky, Gabriel Balmus, Huu Phuc Nguyen, Mahmoud A. Pouladi, Darren G Monckton, Verónica Brito, Paolo Beuzer

**Affiliations:** Departament de Biomedicina, Facultat de Medicina i Ciències de la Salut, Institut de Neurociències, Universitat de Barcelona, Barcelona, Spain; Institut d’Investigacions Biomèdiques August Pi I Sunyer (IDIBAPS), Barcelona, Spain; Centro de Investigación Biomédica en Red Sobre Enfermedades Neurodegenerativas (CIBERNED), Instituto de Salud Carlos III, Madrid, Spain; CHDI Management, Inc., the company that manages the scientific activities of CHDI Foundation Inc., Princeton, NJ 08540, USA; Evotec SE. Manfred Eigen Campus. Essener Bogen 7. 22419 Hamburg, Germany; Movement Disorders Unit, Sant Pau Hospital, Barcelona, Spain; Department of Medicine, Universitat Autònoma de Barcelona (UAB), Barcelona, Spain; Institut de Recerca Sant Pau (IR SANT PAU), Barcelona, Spain; School of Molecular Biosciences, College of Medical, Veterinary and Life Scinces, University of Glasgow, Glasgow, UK; Department of Medical Genetics, Centre for Molecular Medicine and Therapeutics, Djavad Mowafaghian Centre for Brain Health, Edwin S.H. Leong Centre for Healthy Aging, University of British Columbia, Vancouver, BC, Canada, British Columbia Children’s Hospital Research Institute, University of British Columbia, Vancouver, V5Z 4H4, Canada; UK Dementia Research Institute, Department of Clinical Neurosciences, University of Cambridge, Cambridge Biomedical Campus, Cambridge, UK Kangning He, Denny Yang, Gabriel Balmus; Department of Molecular Neuroscience, Transylvanian Institute of Neuroscience, Cluj-Napoca, Romania; Department of Human Genetics, Medical Faculty, Ruhr University Bochum, Bochum, Germany; Servei de Neurologia and Centre d’Esclerosi Múltiple de Catalunya (Cemcat), Institut de Recerca Vall d’Hebron (VHIR), Hospital Universitari Vall d’Hebron, Universitat Autònoma de Barcelona, Barcelona, Spain; Ionis Pharmaceuticals, Inc. 2855 Gazelle Ct. Carlsbad, CA USA 92010; Center for Genomic Medicine, Massachusetts General Hospital, Boston, MA, 02114, USA; Department of Neurology, Harvard Medical School, Boston, MA, 02115, USA; Medical and Population Genetics Program, the Broad Institute of M.I.T. and Harvard, Cambridge, MA, 02142, USA; Institute of Biomedical Research of Barcelona, IIBB-CSIC, Barcelona, Spain

## Abstract

Somatic instability (SI) of expanded DNA repeats is a hallmark of repeat expansion disorder (REDs) and drives onset and progression in Huntington’s disease (HD) yet the absence of target engagement (TE) biomarkers for SI-modulating therapies represents a critical gap to clinical development. Here, we describe the development of the unscheduled repair synthesis assay (URSA)—combining 5-ethynyl-2’-deoxyuridine (EdU) pulse-labeling with digital PCR or sequencing—and show that the CAG-expanded huntingtin (*HTT*) exon 1 allele is a highly active DNA repair hotspot in cells from people with HD (PwHD). Repair activity increases with repeat length, is allele-specific, and depends strongly on MSH3, a central driver of somatic expansion. URSA robustly quantifies MSH3 modulation in preclinical models within days compared to weeks or months required by conventional repeat-length measurements. Critically, substantial repair activity is detectable in peripheral blood mononuclear cells (PBMCs) from PwHD, where signal correlates with CAG length and improves predictive models of somatic expansion (SE) beyond age and CAG length alone. Unlike repeat-length changes, which require years to accumulate in blood, URSA signal is measurable within days. These findings establish DNA repair activity at the mutant *HTT* locus as a mechanistically grounded pharmacodynamic biomarker, enabling TE monitoring on a clinically actionable timescale, and with broad applicability to REDs and other diseases where modulation of the DNA damage response is therapeutically targeted.

## MAIN

Simple DNA repeat expansions beyond a gene-specific threshold underlie several devastating neurological disorders^1–4^. Although historically estimated to affect ∼1 in 3,000 individuals worldwide, recent genomic studies suggest pathogenic repeat expansions causing neurological diseases may occur in as many as 1 in 283 individuals^5^. Huntington’s disease (HD), an archetypal repeat expansion disorder (RED) is caused by the inheritance of 36 or more CAG repeats in exon 1 of the huntingtin (*HTT*) gene, with expansions ≥ 40 being fully penetrant. Beyond the inherited mutation, somatic expansions of the CAG-repeat tract accumulate over time, particularly in vulnerable striatal neurons, which has been shown to drive disease onset, progression, and neurodegeneration^6–9^. Given the central role of somatic instability (SI) in disease progression, considerable effort has focused on identifying the molecular pathways that drive this process. Genome-wide association studies (GWAS) in people with HD (PwHD) have identified DNA repair and handling genes, particularly components of DNA mismatch repair (MMR) pathway as genetic modifiers of disease severity^10^ and somatic expansion (SE) of the *HTT* CAG repeat in blood^11^. Consistent with this, genetic and pharmacological modulation of MMR factors suppresses CAG-repeat instability in HD mouse models and PwHD-derived cells^6,12–16^. Together, these findings establish MMR-driven SI as a central disease mechanism and a compelling therapeutic target. Among these factors, MSH3 has emerged as a particularly attractive target given its absolute requirement for somatic expansion^6,13,15–19^ and relatively favourable safety profile^6,20–22^. Consequently, several academic and pharmaceutical programs are now actively pursuing therapeutic strategies to modulate SI through the MMR pathway in HD and REDs 13,15,16,23–28.

A major barrier to the clinical development of such therapies is the lack of suitable target engagement (TE) and pharmacodynamic biomarkers. Currently, MMR activity in HD has been inferred by quantifying the accumulation of CAG repeat length changes in blood DNA with age^29^. Although *HTT* SI in blood DNA has been proposed as the most promising clinical biomarker for clinical trials, CAG-repeat length expansions accumulate very slowly in blood^11,30^. Preclinical evaluation of SI in human-derived neurons is also challenging, as CAG repeat expansions can be detected only after several weeks in culture and in cellular models carrying pathogenic alleles within the juvenile-onset HD range which typically harbor very large (>60) CAG-repeat expansions^1,13,15,27,31–33^. To overcome these limitations, we explored whether capturing the DNA repair activity driving SI could provide a more sensitive measure than longitudinal repeat-length changes. Although MMR engagement at CAG-repeat tracts has been proposed ^28,34–37^, there is currently no quantitative assessment of how frequently MMR engages expanded repeat tracts in living cells. We hypothesised that the expanded CAG-repeat tract in *HTT* exon 1 undergoes frequent repair events, many of which would cause no change to repeat length. If so, directly quantifying repair-associated DNA synthesis should provide a sensitive readout of ongoing repair activity. We therefore developed unscheduled repair synthesis assay (URSA), to quantify DNA repair activity at the *HTT* locus. URSA exploits unscheduled DNA synthesis as a sensitive, high-resolution proxy for repair activity^38–41^. We rapidly and sensitively measure unscheduled DNA synthesis in a fast and sensitive manner by pulse-labelling non-dividing cells with the thymidine analogue 5-ethynyl-2′-deoxyuridine (EdU) and quantifying its incorporation by sequence capture using either digital PCR (dPCR) or sequencing. Using URSA, we show for the first time that DNA repair activity at the endogenous expanded *HTT* allele is readily measurable in both human induced pluripotent stem cells (iPSCs)-derived neurons and *ex vivo* cultured human HD peripheral blood mononuclear cells (PBMCs). Repair occurs more frequently than previously appreciated, establishing the *HTT* locus as a DNA repair hotspot in HD cells. The repair activity at the *HTT* locus correlates with CAG-repeat length and SI and is dependent on MSH3. Therefore, DNA repair activity at the *HTT* locus represents a promising pharmacodynamic and TE biomarker to support the development of therapeutics targeting MMR and SE.

## RESULTS

### Genome-wide mapping identifies *HTT* exon 1 as a DNA repair hotspot in HD stem cell derived neurons

Expanded CAG repeats are hypothesized to form secondary DNA structures^28,42–44^, that are recognized by the MMR machinery^45–48^. Although MMR proteins can engage mispaired CAG repeats and influence repeat instability, the frequency of this engagement *in vivo,* and whether this region exhibits elevated repair activity relative to the rest of the genome remains unknown. To address this, we measured DNA repair activity at the mutant *HTT* exon 1 locus in neurons using URSA-seq, an adaptation of Repair-seq described by Reid *et* al^41^. Neurons were differentiated from an allelic series of engineered RUES2 human embryonic stem cells (hESCs) (CAG20, CAG56, CAG72 and CAG118)^49^, H9 hESCs (CAG20 and CAG81)^50^ and an HD patient-derived iPSC carrying a CAG125 allele^15^. No dividing cells were detected (Extended Data 1a, b), confirming that any EdU incorporation reflects repair-associated DNA synthesis rather than replication-associated DNA synthesis. EdU labelling for 3 or 7 days did not affect neuronal viability or global DNA damage responses in any RUES2 genotype tested, as assessed by γH2AX staining (Extended Data 1c-e). After labeling, EdU-positive DNA fragments were enriched and subjected to next-generation sequencing (URSA-seq). Genome-wide peak calling revealed between 60,000 and 150,000 peaks with significant EdU enrichment relative to depth-matched whole-genome sequencing across all cell lines analyzed (Extended Data 2), consistent with previous studies. Peaks were prominently enriched at promoters, 5′-UTRs, gene bodies, and distal intergenic regions at both 3 and 7 days (Extended Data 3a) in agreement with previously reported DNA repair hotspot profiles^40,41^. A substantial proportion of peaks were shared across genotypes and labeling conditions, indicating that EdU incorporation profiles are largely reproducible (Extended Data 3b).

We next examined whether the *HTT* locus showed evidence of repair-associated DNA synthesis. No significant repair hotspots were detected at the endogenous *HTT* locus in RUES2 neurons carrying a non-expanded CAG20 allele at either 3 (Extended Data 3c) or 7 days of EdU treatment (Fig. 1a) indicating minimal baseline repair-associated synthesis at the non-expanded *HTT* locus. In contrast, all RUES2-HD neurons exhibited a prominent and reproducible peak at the *HTT* exon 1. (Fig. 1a, and Extended Data 3c). Peak intensity increased with CAG repeat length at both labeling time points (Fig. 1b and Extended Data 3d). Differential enrichment analysis comparing expanded CAG genotypes to CAG20 identified thousands of loci with significant EdU incorporation differences (False Discovery Rate (FDR) < 0.05). *HTT* was among the most enriched and significant loci in HD cells and showed an increase in effect size with repeat length (Fig. 1c). While *HTT* exon 1 peak in cells not carrying expansions was below the average peak intensity, it was among the highest signal peaks in all HD cell lines (Fig. 1d and Extended Data 3e). The correlation between URSA and CAG length was specific to *HTT* exon 1: neither an intragenic DNA repair hotspot (DRH) (*HTT* exon 32) nor other reference DRHs showed increased signal as a function of CAG length (Fig. 1e, f and Extended Data 3f-g). To determine whether this enrichment generalizes to other HD neuronal models, we examined URSA-seq profiles in H9-derived CAG81 neurons. Consistent with RUES2 HD neurons, *HTT* exon 1 exhibited a prominent peak (Extended Data 4a). Differential analysis confirmed significant enrichment relative to H9 controls (Extended Data 4b) whereas signals across other DRHs remained comparable (Extended Data 4c-e). A clear *HTT* exon1 peak was also observed in patient-derived, non-engineered iPSC neurons with a CAG125 allele (Extended Data 5a). Notably, *HTT* exon 1 ranked consistently in the top 1% of genome-wide DRHs across all HD lines, including CAG56 (Supplementary Table 1). Together, these data establish that expanded CAG repeats are sufficient to drive focal, repeat length-dependent repair activity at *HTT* exon 1, identifying this region as a prominent DNA repair hotspot in HD neurons.

**Main Figure 1:**
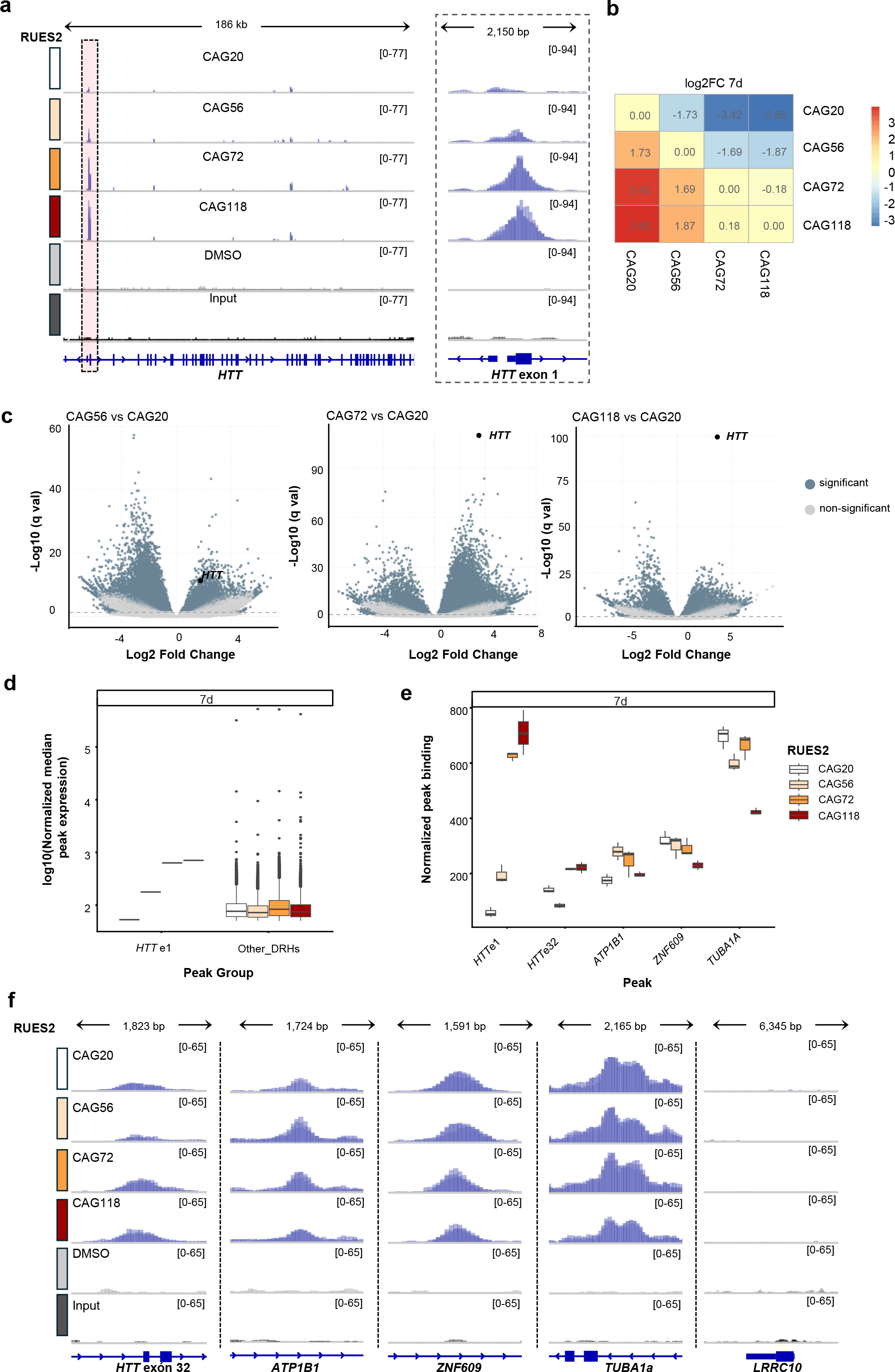
Genome-wide URSA-seq profiling reveals substantial unscheduled repair synthesis at the *HTT* exon 1 locus in an allelic series of RUES2 derived neurons (a) Integrative Genomics Viewer (IGV) tracks at the *HTT* locus following 7 days of EdU treatment (20 µM) in RUES2 neurons (CAG20, CAG56, CAG72, and CAG118). DMSO-treated samples (negative control) and input DNA are shown alongside the enriched libraries. A focal EdU enrichment peak is present at *HTT* exon 1 (red box, zoomed right) in HD genotypes but not in controls. N=3wells from one neuronal differentiation **(b)** Heatmap of log2 fold-change (Log2FC) in EdU-enrichment at 7 days normalized to input, across genotypes. **(c)** Volcano plots of differential EdU enrichment comparing HD neurons (CAG56, CAG72 and CAG118) with WT (CAG20). Significant peaks (False Discovery Rate (FDR) < 0.05, |log₂FC| > 1) are shown in green. The *HTT* exon 1 DNA repair hotspot (DRH) is indicated and exhibits progressively greater enrichment with increasing CAG repeat length. **(d)** Comparison of normalized *HTT* exon 1 peak (*HTT*exon1) intensities against all other DRHs peaks (other_DRHs), by genotype after 7 days EdU incorporation. DRHs were defined as peaks called in at least two biological replicates per genotype, with ≥50 normalized counts, and absent in the DMSO control. *HTT* exon 1 signal is below median DRH intensity in WT neurons and among the highest in all HD genotypes. **(e)** Normalized *HTT* exon 1 signal relative to other DRHs loci, by genotype. While *HTT* exon1 peak height increases with longer CAG length, other DRHs remain stable. Error bars indicate mean ± s.e.m. **(f)** IGV tracks at representative reference loci with consistent signal across genotypes. N=3 wells from one neuronal differentiation

### Quantitative locus-specific measurement of DNA repair at *HTT* exon 1

To quantify repair activity at the *HTT* locus with greater sensitivity and throughput, we developed URSA-dPCR, where targeted digital PCR on EdU-enriched DNA fractions is used in place of next generation sequencing, enabling locus-specific quantification of repair activity (Fig. 2a). Both non-allele-specific and allele-specific primers were designed in the 5′ flanking region upstream of the *HTT* CAG tract to ensure consistent amplification across alleles of different repeat lengths. Allele specificity was achieved using a previously described informative SNP^49^ (Fig. 2b). Reference loci were selected from stable DRHs identified in both our URSA-seq (Fig. 1f and Extended Data 3g, 4e) and published datasets^40,41^. URSA-dPCR detected a strong signal in RUES2 CAG72 at the *HTT* exon 1 locus with both non-allele specific and expanded allele-specific primers. In contrast, non-expanded allele specific primers detected a very low signal - consistent with basal repair activity near transcription start sites^41^, demonstrating that the expanded allele accounts for the vast majority of the *HTT* exon 1 signal (Fig. 2c). Signal at reference loci *HTT*e32, *ATP1B1*, and *ZNF609* remained comparable between genotypes, while no signal was detected at the negative control locus *LRRC10* (Fig. 2c).

**Main Figure 2:**
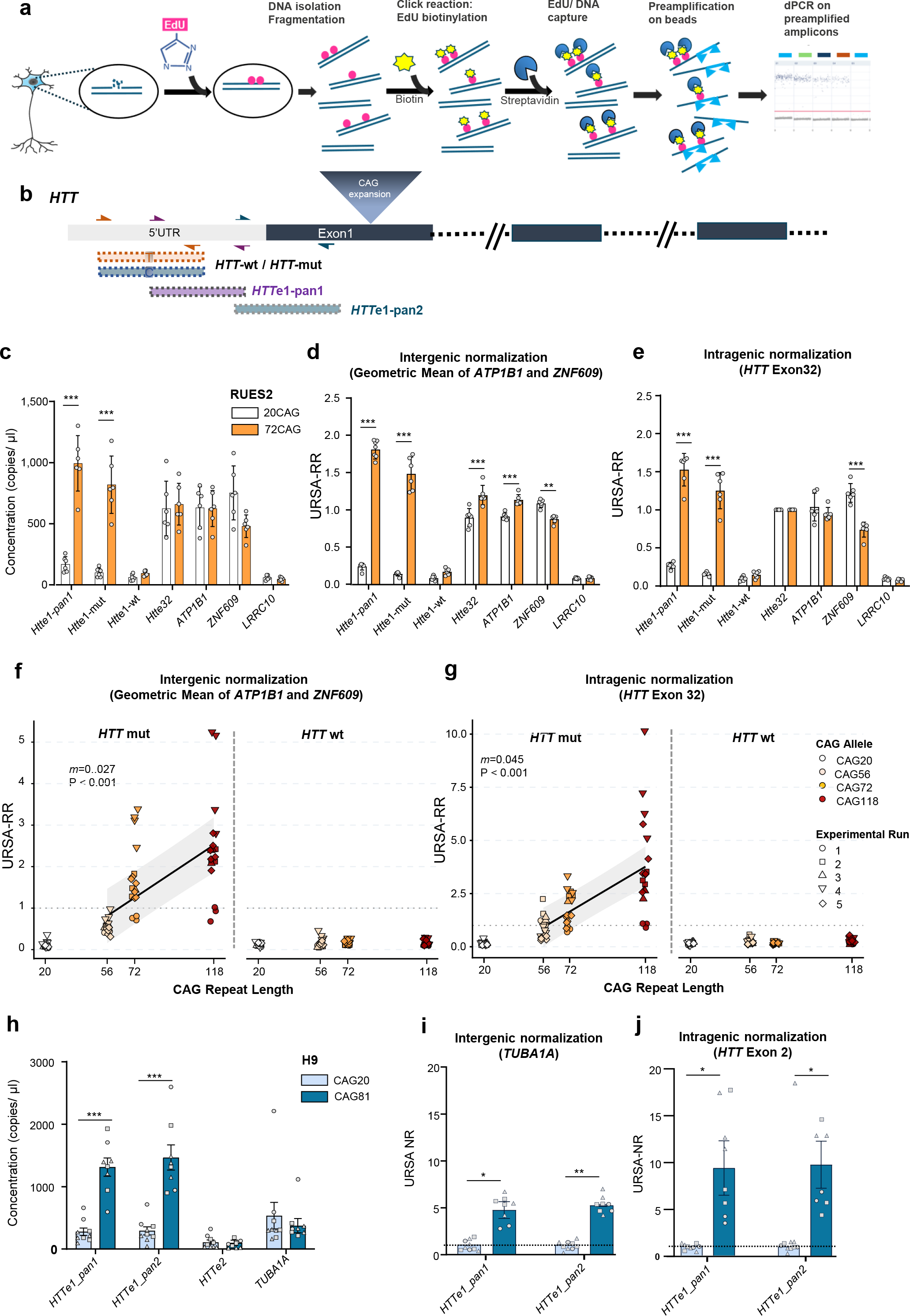
*HTT* exon 1 repair activity is sensitively quantified by URSA-dPCR in ESC-derived neuronal lines. **(a)** Overview of the URSA-dPCR workflow. Neurons were incubated with EdU for 3 or 7 days. DNA was extracted and fragmented, EdU-labelled DNA fragments were biotinylated by click chemistry, purified with streptavidin beads, and quantified at loci of choice by digital PCR (dPCR or ddPCR). **(b)** Schematic of the *HTT* locus showing primer pairs targeting exon 1 and the 5′ UTR SNP rs146151652, located upstream of the CAG repeat, used for allele-specific detection. **(c)** URSA-dPCR raw signal (copies/µL) at *HTT* exon 1 (*HTT*e1-pan1), the expanded (*HTT*mut), and non-expanded (*HTT*wt) alleles, *HTT* exon 32, *TUBA1*, *ATP1B1*, *ZNF609* and *LRRC10* in wild type RUES2-CAG20 or HD RUES2-CAG72 neurons. ***P < 0.001, unpaired two-tailed t-test. N=6 experimental runs, i.e., number of independent neuronal differentiations **(d)** URSA-dPCR signal normalized to the geometric mean of two stable reference loci (*ATP1B1* and *ZNF609*) (URSA-RR). RUES2-CAG72 neurons exhibit a marked signal increase at mutant *HTT* exon 1 relative to RUES2-CAG20, whereas reference loci remain similar between genotypes. ***P < 0.001, unpaired two-tailed t-test. N=6 experimental runs, i.e., number of independent neuronal differentiations **(e)** As in **(d)**, normalized to *HTT* exon 32 (URSA-RR). ***P < 0.001, unpaired two-tailed t-test. N=6 experimental runs, i.e., number of independent neuronal differentiations **(f, g)** Allele-specific *HTT* exon 1 repair signal (URSA-RR) at the mutant (left) and wild-type (right) alleles across the CAG allelic series after 7 days of EdU treatment. URSA-RR was normalized to the geometric mean of *ATP1B1* and *ZNF609* **(f)** or to *HTT* exon 32 **(g)**. Each point represents one well; shape denotes the independent neuronal differentiation (N = 5) with 2–6 wells per differentiation. Solid lines in the mutant panels represent linear mixed-effects models with CAG repeat length as a fixed effect and neuronal differentiation as a random intercept (f: *slope = m* = 0.027, P = 7.42 × 10⁻¹¹; **g**: *m* = 0.04, P = 3.4 x 10^-09^). **(h)** URSA-ddPCR raw signal (copies/µL) at *HTT* exon1 using 2 sets of primers (*HTT*e1-pan1 and *HTT*e1-pan2), *HTT* exon 2, and *TUBA1A* in H9 CAG20 and CAG81 neurons after 7 days of EdU incorporation. Bars represent the mean ± s.e.m. of three independent neuronal differentiations. Individual points represent single wells (2–4 wells per differentiation), with symbol shape indicating the independent neuronal differentiation. ***P < 0.001, two-way ANOVA. **(i, j)** Normalized raw URSA signal (URSA-NR) at the *HTT* exon 1 DNA repair hotspot (DRH) in H9-derived neurons using **(i)** intergenic normalization to *TUBA1A* and **(j)** intragenic normalization to *HTT* exon 2 for both primer sets *(HTT*e1-pan1 and *HTT*e1-pan2). For graphical representation, *HTT* exon 1 signal was normalized to the indicated reference and scaled to the H9 CAG20 baseline. Statistical analyses were performed on the corresponding unscaled raw normalization ratios (*HTT*e1/reference). Bars represent the mean ± s.e.m. of three independent neuronal differentiations. Individual points represent single wells 2–4 wells per differentiation, with symbol shape indicating the independent neuronal differentiation. *P < 0.05, **P < 0.01, paired two-tailed *t*-test. Increased EdU incorporation at *HTT* exon 1 is observed in all expanded genotypes, consistent with enhanced repair activity at the *HTT* exon 1 DRH.

To accurately quantify repair activity at the *HTT* locus and enable comparison across experiments, the m*HTT* signal was normalized to stable reference regions. Both intergenic normalization to the geometric mean of *ATP1B1* and *ZNF609*, and intragenic normalization to *HTT* exon 32 demonstrated elevat*ed HTT* exon 1 URSA-dPCR signal (Fig. 2d-e). We next extended this analysis across the full RUES2 isogenic series to determine whether URSA signal scales with CAG length. Both normalization strategies revealed a highly significant positive association between URSA-dPCR signal at the mutant *HTT* allele and inherited CAG length at both 3 and 7 days of EdU incorporation (p < 0.001; Fig. 2f, g and Extended Data 3h, i). Notably, the x-intercept of the regression line fell at approximately 26.5 CAG repeats — below the intermediate allele threshold of 36 — suggesting that detectable repair activity at the *HTT* locus may begin to accumulate prior to the established pathological range. This is consistent with reports of low-level SI at inherited CAG lengths below 36^51,52^. Elevated repair activity at *HTT* exon 1 was not restricted to the RUES2 isogenic series. H9-derived neurons carrying an expanded CAG81 allele also exhibited significantly increased *HTT* exon 1 URSA-dPCR signal compared with CAG20 controls under both intergenic and intragenic normalization strategies (Fig. 2h-j). Furthermore, patient-derived, non-gene-edited CAG125 iPSC neurons displayed URSA-dPCR signals comparable to those observed in RUES2 CAG118 neurons (Extended Data Fig. 5). Together, these findings establish that the expanded *HTT* exon 1 is a highly engaged DNA repair hotspot in HD neurons and demonstrate that URSA-dPCR sensitively quantifies repair activity at this locus across independent neuronal models.

### MSH3 drives the high repair activity at the mutant *HTT* hotspot revealed by URSA

*MSH3* is one of the strongest and best established genetic modifiers of both CAG repeat expansion and disease onset in HD^11,53–56^. MSH3 and MSH2 form the heterodimer MutSβ which plays a central role in repeat somatic expansion by recognizing extrahelical extrusions at CAG repeats^28,57^. To test whether the URSA signal at *HTT* exon 1 depends on MSH3, we modulated MSH3 levels using multiple, independent approaches. We first generated *MSH3* knock-out (KO) clones of RUES2 CAG118 ESCs, differentiated them in neurons, and confirmed protein loss (Fig. 3a). URSA-dPCR and URSA-seq analysis of two independent MSH3 KO clones (Cl10 and Cl72) showed a greater than 90% reduction in signal at *HTT* exon 1, with no significant change at the reference loci *ATP1B1*, *ZNF609*, *LRRC10*, and *HTT* exon 32 (Fig 3b-d). This reduction was striking with both intergenic (*ATP1B1*, *ZNF609*) and intragenic (*HTT* exon 32) normalization and was further confirmed by URSA-seq, in which the *HTT* exon 1 peak was comparable to CAG20 levels in both KO clones (Fig 3d).

**Main Figure 3:**
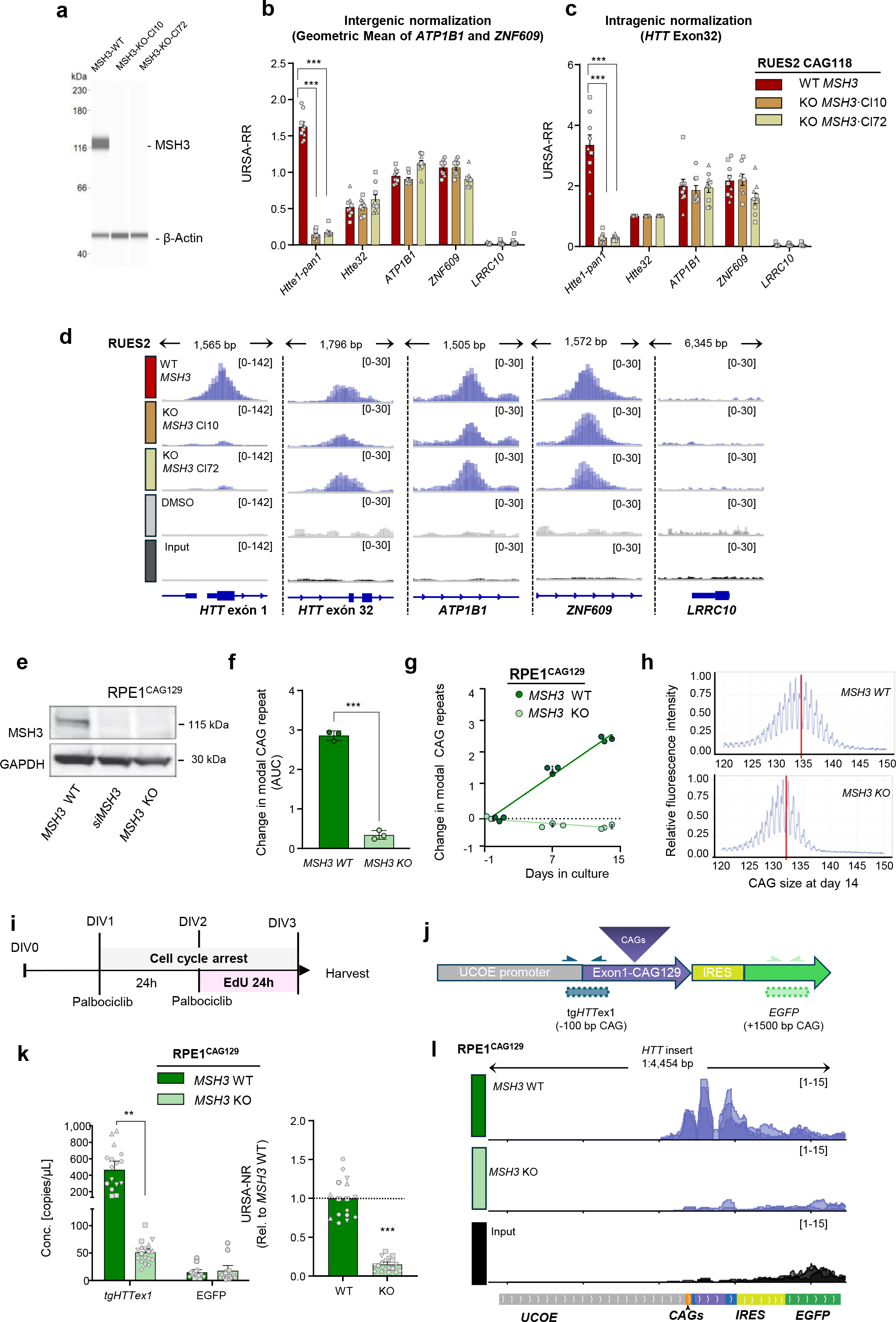
Repair-associated EdU incorporation at *HTT* exon 1 is MSH3-dependent. **(a)** Representative Western Blot of MSH3 confirming MSH3 knockout in clones CL10 and CL72. Bands show MSH3 (∼115 kDa) and β-actin, (∼42 kDa) as loading control. **(b, d)** Normalized URSA-dPCR signal (URSA-RR) in RUES2 CAG118 and two RUES2 CAG118 *MSH3* KO clones (CL10, CL72). Signal at *HTT* exon 1 is significantly reduced in MSH3-KO clones, while signal at reference loci is unaffected, under **(b)** intergenic and **(c)** intragenic normalization. Bars represent the mean ± s.e.m. of 3 independent neuronal differentiations. Individual points represent one well (2–4 wells per differentiation), with symbol shape indicating the independent neuronal differentiation. **P < 0.001, paired two-tailed t-test **(d)** IGV genomic tracks of URSA-seq signal at *HTT* exon 1 and reference DRH loci in RUES2 CAG118 and *MSH3* KO clones CL10 and CL72. The *HTT* exon 1 peak is nearly abolished in both KO clones, while signal at reference loci remains unchanged. N=3 wells from one neuronal differentiation **(e)** Western blot of MSH3 in RPE-1^CAG129^ cells: parental (*MSH3* WT), siRNA-depleted (siMSH3), and *MSH3* KO clone. GAPDH, loading control. Bands show MSH3 (∼115 kDa) and GAPDH (∼30 kDa) as loading control. **(f-h)** Somatic expansion of the CAG tract in RPE-1^CAG129^ *MSH3* WT and *MSH3* KO cells over 14 days, measured by capillary electrophoresis. **(f)** Change in modal CAG repeat length relative to day 0 ***P < 0.001, unpaired two-tailed t-test **(g)** AUC quantification of expansion across the time course. Data are mean ± s.d.; N = 3 independent experimental runs per genotype. **(h)** Representative capillary electrophoresis traces at day 14 in *MSH3* WT (top) and *MSH3* KO (bottom) cells. Red vertical line, modal CAG length. The *MSH3* WT distribution is shifted toward longer repeats relative to *MSH3* KO. **(i)** URSA labelling scheme in RPE-1^CAG129^ cells: cultures were seeded at DIV0, arrested in G1 by CDK4/6 inhibition (palbociclib; 250nM) at DIV1 and DIV2 (24 h pre-treatment), pulsed with EdU for 24 h from DIV2 to DIV3, and harvested at DIV3. **(j)** schematic of the integrated *HTT* exon 1 CAG129 transgene: a UCOE promoter drives *HTT* exon 1 with 129 CAGs, followed by an IRES and *EGFP*. Arrowheads indicate dPCR amplicon positions: tg*HTT*ex1 (–100 bp upstream of the CAG tract) and *EGFP* (+1,500 bp). **(k)** Left: URSA-dPCR quantification (copies**/**μl) at tg*HTT*ex1 and *EGFP* in RPE-1^CAG129^ MSH3 WT and MSH3 KO cells; N = 6 independent experimental runs per genotype. Bars represent the mean ± s.e.m. Individual points represent one culture (2–4 cultures per run), with symbol shape indicating the independent runs. \*\**P* < 0.01, paired two-tailed *t*-test. Right: URSA normalized ratio (URSA-NR; tg*HTT*ex1*/EGFP*) relative to the mean of the corresponding RPE-1^CAG129^ MSH3 WT cells from each independent run (defined as 1; dashed line). Bars represent the mean ± s.e.m. of N=6 independent experimental runs. Individual points represent single wells (2–4 wells per run) with symbol shape indicating the independent runs. \*\*\**P* < 0.001, one-sample two-tailed *t*-test against the null value of 1 (*t*(5) = −29.35). **(l)** IGV tracks of URSA-seq signal at *HTT* exon 1 in RPE-1^CAG129^ *MSH3* WT, *MSH3* KO cells, and input. The *HTT* exon 1 peak is nearly abolished in the KO cells. N=3 wells from one experimental run

To determine whether these findings generalize beyond neuronal cell lines, we examined RPE-1 cells stably expressing an *HTT* exon 1 construct with 129 CAG repeats^33^. As expected, CRISPR-Cas9 deletion of *MSH3* (Fig. 3e) abolished CAG somatic expansion (Fig. 3f-h). To perform URSA in these proliferating cells, cultures were arrested in G1 with palbociclib prior to a 24-hour EdU pulse **(Fig. 3i);** successful cell cycle arrest was confirmed by flow cytometry (Extended Data 6a, b). *HTT* exon 1 URSA-dPCR signal was normalized to the EGFP cassette within the same construct and expressed relative to WT cells (URSA-NR) (Fig. 3j, k). G1-arrested RPE1^CAG129^ cells produced a robust URSA signal at the transgenic *HTT* locus, which was reduced by approximately 85% upon *MSH3* KO as assessed by both URSA-dPCR (Fig. 3k) and URSA-seq (Fig. 3l).

To evaluate whether therapeutically relevant *MSH3* reduction strategies produce comparable effects, we treated neurons with siRNA or AAV-delivered shRNA targeting *MSH3* (Fig. 4a, Extended Data 7a). Both approaches achieved greater than 80% MSH3 reduction (Fig. 4b, c; Extended Data 7b, c) and caused greater than 80% reduction in URSA-dPCR signal at *HTT* exon 1 without significantly altering repair activity at reference loci (Fig. 4d, e; Extended Data 7d-f). In summary, *MSH3* depletion by all three modalities — genetic KO, siRNA, and AAV-shRNA — reduced repair-associated DNA synthesis at the *HTT* locus without impacting repair activity at other reference DRHs (Fig. 3b, c; Fig. 4d, e; Extended Data 7d, e). Taken together, these results establish that the elevated DNA repair synthesis detected by URSA at mutant *HTT* exon 1 is MSH3-dependent and parallels CAG somatic expansion, directly linking the repair hotspot to the MutSβ pathway that drives instability and modifies HD onset and progression.

**Main Figure. 4:**
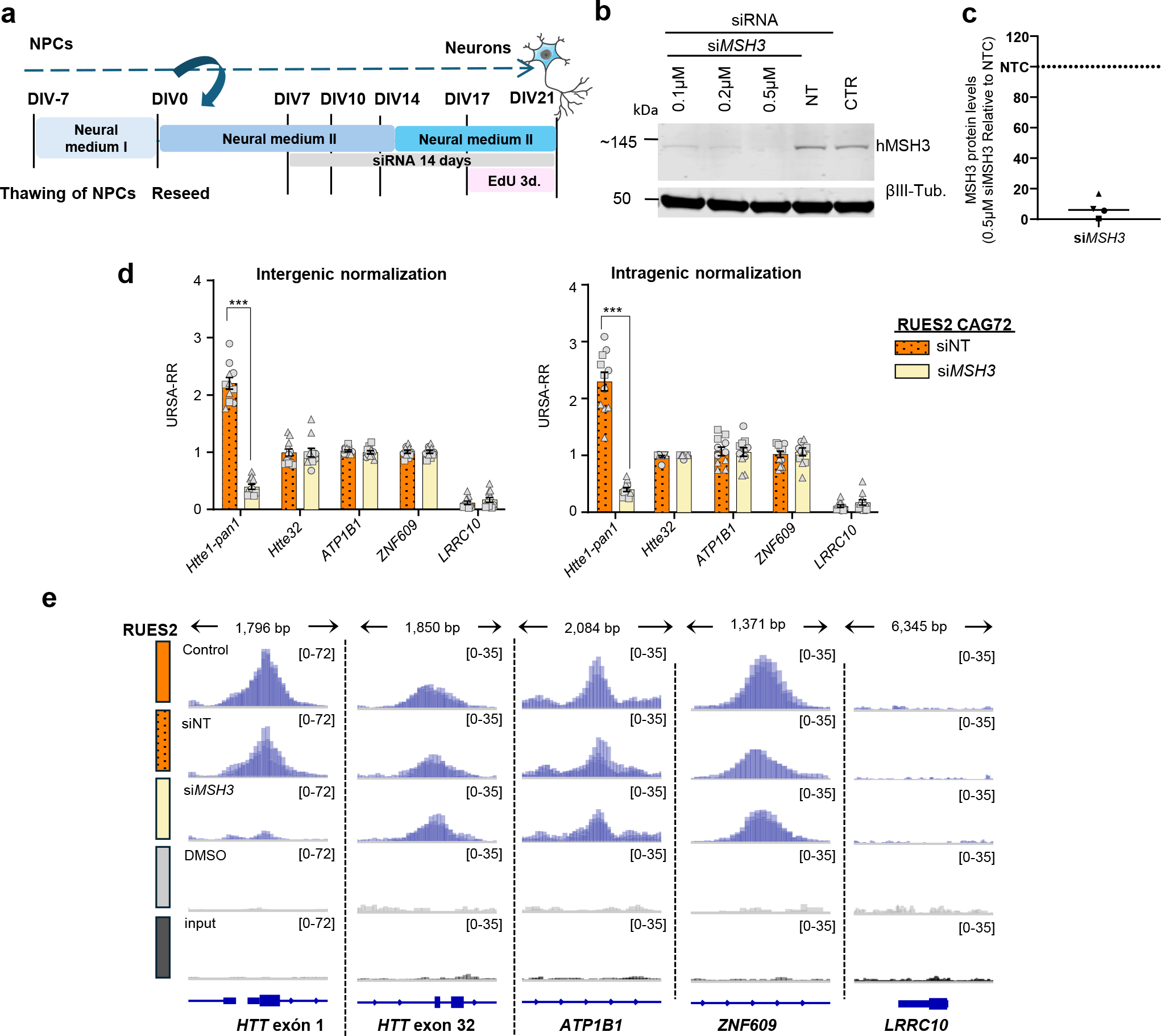
URSA detects MSH3 modulation by therapeutically relevant approaches in RUES2 derived neurons. (a) Schematic of neuronal differentiation, siRNA transfection timing and EdU labeling in RUES2-72CAG neurons. Cells were transfected with siRNA at the indicated time point and exposed to EdU for 3–4 days within the defined labeling window prior to sample collection. (b) Representative Western Blot confirming MSH3 knockdown by siRNA in CAG72 neurons. Bands show MSH3 (∼115 kDa) and βIII-tubulin (55 kDa) as loading control. (c) Quantification shows a robust reduction in MSH3 protein levels following siRNA treatment. (d) Intergenic (left) and intragenic (right) normalized URSA-dPCR signal at the HTT exon 1 locus after transfection with non-targeting or MSH3 targeting siRNA. Values were scaled to the non-targeting siRNA condition (URSA-RR); only 18–19% of HTT exon 1 URSA-dPCR signal remains after MSH3 knock-down. Bars represent the mean ± s.e.m. of three independent neuronal differentiations. Individual points represent one culture (3–4 wells per differentiation), with symbol shape indicating the independent neuronal differentiation (N=3). ***P < 0.001 paired two-tailed t-test. (e) IGV tracks of URSA-seq signal at HTT exon 1 and representative reference loci (HTT exon 32, ATP1B1, ZNF609, LRRC10) in CAG72 RUES2 neurons treated with MSH3 siRNA or non-targeting siRNA (siNT) for 3 days compared with DMSO and input controls. Stable loci show consistent signal across conditions, while the HTT exon 1 repair hotspot is markedly reduced upon MSH3 depletion. N=3 wells from one neuronal differentiation

### Detection of a DNA repair hotspot at the *HTT* locus in HD PBMCs using URSA digital PCR

Given their accessibility as a clinically relevant tissue, we asked whether repair activity at *HTT* exon 1 could be detected in PBMCs. We applied URSA to PBMCs from 31 individuals with HD-causing CAG expansions and 25 unaffected controls (Fig. 5a; Supplementary Table 2), spanning intermediate (35 CAG, n=1), reduced-penetrance (36–39 CAG, n=1), full-penetrance (40–54 CAG, n=25) and juvenile-onset (≥55 CAG, n=5) alleles. Fluorescent Activatd Cell Sorted analysis confirmed negligible proliferation after 24 h of EdU treatment (Extended Data Fig. 8), indicating that EdU incorporation reflected repair-associated synthesis rather than DNA replication.

**Main Figure 5:**
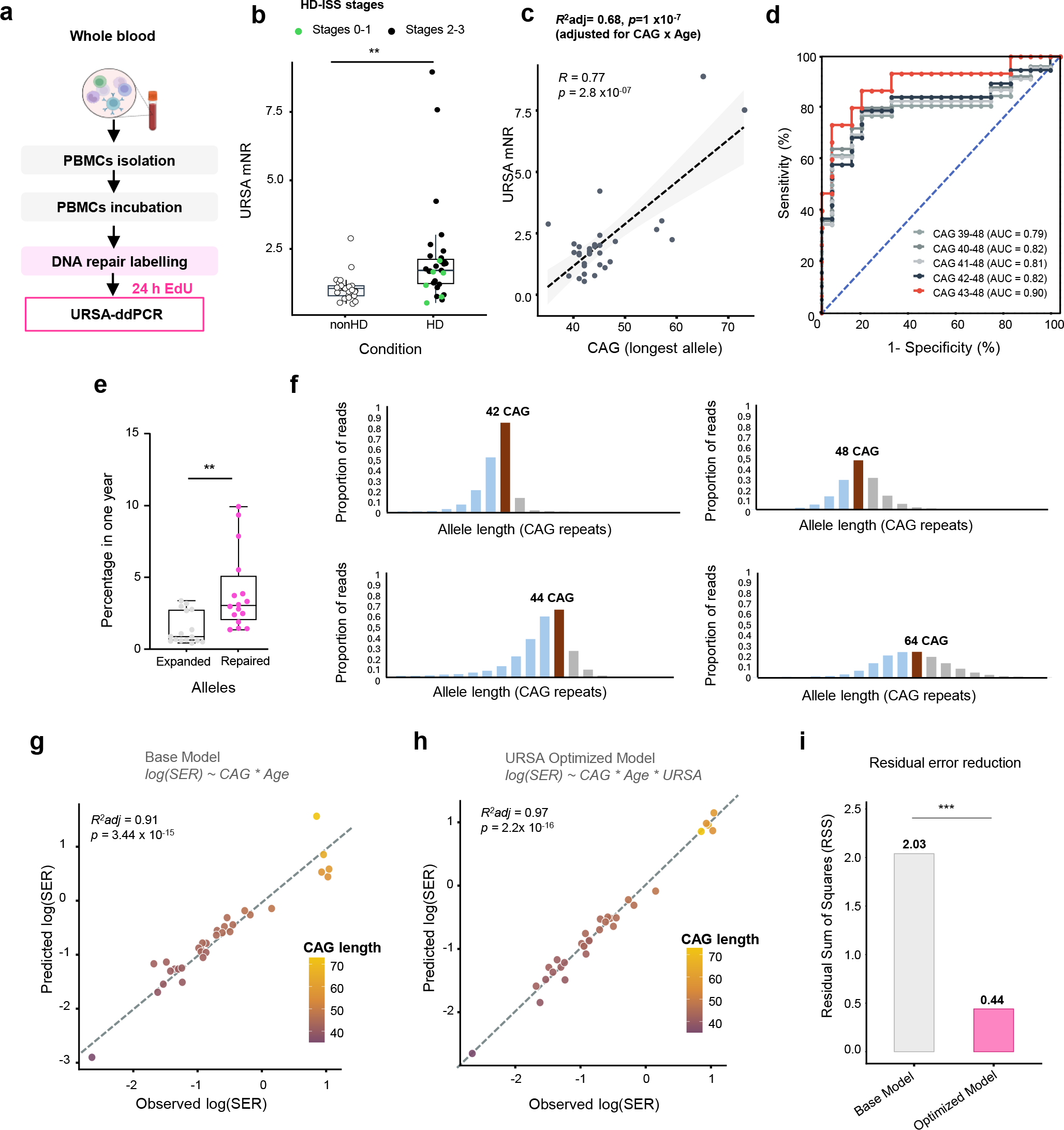
Repair activity at the *HTT* exon 1 hotspot predicts CAG-repeat instability in peripheral blood mononuclear cells. **(a)** Schematic representation of the experimental workflow for PBMC analysis: whole blood collection was followed by PBMC isolation (2 h) and a brief stabilization period (2 h) prior to EdU addition. PBMCs were then incubated with EdU (20 µM) to label DNA repair synthesis for 24 h, after which samples were processed for URSA-dPCR analysis. **(b)** URSA-mNR in Non-HD individuals and *HTT* expansion carriers. *HTT* expansion carriers are colour-coded according to clinical status (individuals in HD-ISS stages 0-1, green; in HD-ISS 2-3, black). The Non-HD group includes one individual carrying a 35-CAG intermediate allele who had neurological symptoms but did not meet diagnostic criteria for HD. Each point represents one donor; boxplots show the median and interquartile range. **P < 0.01, unpaired two-tailed t-test **(c)** Relationship between URSA-mNR and expanded *HTT* CAG repeat length. Data include individuals carrying intermediate (35 CAG), reduced penetrance (36–39 CAG), full penetrance (40–54 CAG), and juvenile-onset (≥55 CAG) alleles. The dashed line represents the linear regression and grey shading the 95% confidence interval. Pearson’s correlation coefficient (R) is shown. The adjusted R² and P value are from the multiple regression model incorporating CAG repeat length, age, and the CAG × age interaction. **(d)** Receiver operating characteristic (ROC) analysis evaluating the performance of URSA-mNR to distinguish *HTT* expansion carriers above different CAG thresholds from unaffected controls. The 43–48 CAG threshold, corresponding to the onset of the adult-onset pathogenic range, yielded the highest diagnostic performance (area under the curve (AUC) = 0.897, sensitivity = 86.7%, specificity = 83.3%). **(e)** Percentage of alleles per year carrying CAG expansions (grey) compared to repair-associated events measured by URSA-dPCR (magenta) across *HTT* alleles with increasing repeat length. Values were derived from repeat-length distributions to estimate the proportion of cells with CAG length changes and the annual probability of CAG gain at *HTT*. Each additional CAG repeat was counted as an independent expansion event (for example, a CAG+4 gain equals four expansion events). Repair frequency was calculated as the percentage of EdU-enriched haploid locus copies recovered over the total genomic DNA input, mathematically back-adjusted for sample-processing volumes and pre-amplification cycles (n =16 HD patients, **P < 0.01, unpaired two-tailed t-test) **(f)** Representative distributions of *HTT* CAG repeat lengths determined by MiSeq analysis for selected alleles (42, 44, 48, and 64 CAG). Histograms show proportional read counts across repeat sizes relative to the total number of expanded allele reads **(g)** Scatter plot illustrating the predicted versus observed log(SER) ratio based for the baseline model incorporating age, CAG length, and their interaction (Age × CAG). **(h)** Predicted versus observed log(SER) values for the optimized model including URSA-mNR in addition to CAG repeat length, age and their interaction. Points are coloured according to inherited CAG repeat length. The dashed line indicates the line of identity (y = x). **(i)** Comparison of the residual sum of squares (RSS) between the baseline (CAG x Age) and the optimized URSA-extended model by ANOVA, demonstrating a significant reduction in unexplained variance upon the inclusion of URSA-mNR (F = 21.79; ANOVA ***P < 0.001)

URSA-dPCR measurements from two *HTT* exon 1 primer sets were normalized to intergenic (*RPP30*) and intragenic (*HTT* exon 2) reference loci and averaged into a per-sample URSA mean normalized ratio (URSA-mNR) (Extended Data 9a, b). URSA-mNR significantly distinguished HD donors from controls (P =0.0056; Fig. 5b). Notably, elevated repair activity at *HTT* exon 1 was detected not only in individuals in HD-ISS stage 2-3 but also in several individuals in stage 0-1, indicating that it was detectable prior to clinical onset. Elevated URSA signal was observed across the spectrum of repeat lengths, including the intermediate allele, reduced-penetrance alleles, and short full penetrance alleles (≤42 CAG) (Fig. 5c). Repair activity increased with expanded CAG length, reaching its highest values in PBMCs from people with large and juvenile-onset expansions, indicating that URSA captures a biological process that scales with repeat size. Consistent with this, URSA-mNR correlated strongly with CAG length within the expansion-carrying cohort (r=0.81, P =2.53×10⁻⁸), and a model incorporating CAG length, age and their interaction explained 68% of the variance in repair activity (adjusted R²=0.68, p=2×10⁻⁷).

To determine the repeat-length range over which URSA-mNR most effectively distinguishes carriers from controls, we performed ROC analyses across progressively narrower CAG windows (Fig. 5d). For the broadest window (39–48 CAG), URSA-mNR yielded an area under the curve (AUC) of 0.796; raising the lower threshold to 40–42 CAG produced only modest gains (AUC 0.806–0.820), whereas restricting to 43–48 CAG substantially improved discrimination (AUC=0.897). Together, these findings demonstrate that URSA-mNR captures a repeat-length-dependent signal detectable before clinical onset, with the clearest carrier–control separation observed among individuals with ≥43 CAG repeats.

To relate URSA-detected repair events to actual CAG length-change events, we used ultra-deep MiSeq sequencing of the *HTT* repeat (Fig. 5d, e). We scaled EdU-labeled *HTT* molecules to an equivalent annual rate, assuming 100% capture and PCR efficiency—a conservative estimate representing a lower bound on true repair activity. Even under these highly conservative assumptions, the inferred annual rate of repair-associated events exceeded the annual rate of CAG-gain events detected by sequencing (Fig. 5f), suggesting that URSA captures frequent repair-associated synthesis that does not necessarily result in repeat length change.

To determine whether the repair signal is associated with somatic expansion, we calculated the somatic expansion ratio (SER) as the ratio of MiSeq reads exceeding the modal repeat length to those at the modal repeat length^11,29^. Consistent with previous studies, SER exhibited the expected dependence on CAG repeat length and age at sampling. This relationship was modeled using a linear regression including CAG repeat length, age at sampling, and their interaction (R² = 0.9075, P = 3.44 x 10^-15^. Fig. 5g). Notably, incorporating URSA-mNR into the model explained substantial additional variation in SER (adjusted R² = 0.972, P = 2.2 × 10^-16^, Fig. 5h). This improvement was driven by a significant reduction in the Residual Sum of Squares (RSS) (Fig. 5i) suggesting that URSA-mNR captures individual-specific variance in somatic expansion rate not accounted for by age and CAG length alone. Leave-one-out cross-validation further demonstrated lower prediction error for the URSA-extended model than for the baseline model (Supplementary Table 3). Together, these results indicate that URSA-dPCR detects repeat length–dependent repair activity at the *HTT* exon 1 hotspot in peripheral cells and reflects individual-specific variation in somatic expansion supporting URSA-mNR as a candidate biomarker that directly reads out CAG-dependent disease biology.

## DISCUSSION

Somatic instability in HD and other REDs has become an attractive central target for therapeutic intervention, yet no pharmacodynamic SI assay currently exists to monitor SI-modulating therapeutics now in development. Despite SI being a key driver of HD onset and progression, the mechanisms by which DNA repair pathways engage pathogenic repeats in human cells remain poorly understood. By analyzing unscheduled DNA repair synthesis, we demonstrate that mutant *HTT* exon 1 becomes a highly active DNA repair hotspot. Repair activity at this locus increases with CAG repeat length, is highly specific to the expanded allele, and strongly depends on MSH3 activity in neurons and other preclinical cellular models. Using URSA, we show that this repair activity can be reliably quantified and serves as a robust readout of MSH3 modulation in preclinical models. Critically, a substantial URSA signal is readily quantifiable in PwHD PBMCs, which have been proposed as a tractable clinical TE model^30^. Although these cells exhibit only modest SI requiring years before significant repeat length changes become quantifiable, the upstream repair activity driving instability is readily measurable within days. Together, these findings establish repair activity at m*HTT* as a promising TE biomarker for SI-modulating therapeutics, addressing a critical unmet need in the development of disease-modifying drugs for HD.

URSA offers a functional readout of DNA repair engagement that bridges the gap between genetic evidence implicating MMR^10,11^ and its actual activity at expanded repeats. This engagement is consistent with the formation of non-canonical DNA structures at expanded CAG repeats that may arise during transcription, repair, and replication^58–60^ and trigger MMR activity. The strong dependence of the URSA signal on MSH3 establishes a direct cellular link between MMR activity and *HTT* CAG-repeat expansion. Loss of MSH3 nearly abolished repair-associated DNA synthesis at *HTT* exon 1 without affecting other reference genomic hotspots, indicating that MutSβ-mediated MMR is a central driver of repair specific to the expanded *HTT* locus. The URSA signal likely reflects repeated MutSβ engagement events that do not necessarily correspond to repeat-length changes. Repair outcomes may depend on the nature and size of the mismatch, which determines repair pathway choice, the length of the resynthesized DNA patch, and the involvement of both MMR and non-MMR proteins such as FAN1^61,62^. Frequent cycles of MMR activation and repair-associated DNA synthesis can create opportunities for repeat length changes, ultimately promoting the gradual somatic expansion observed in HD tissues^7–9,57,63,64^. Determining whether, and if so how additional repair components or pathways contribute to synthesis events detected by URSA will require further investigation.

URSA fundamentally changes how MMR modulation can be studied in preclinical settings, delivering functional readouts in days rather than weeks, at scale, and with a sensitivity that existing approaches cannot match. This will potentially enable genetic screens, structure–activity relationship studies, and validation of drug candidates. In principle, URSA could also be adapted to *in vivo* assessment of SI modulators in animal models in both central and peripheral tissues.

Importantly, our study demonstrates that the *HTT* DNA repair hotspot can be interrogated by URSA-dPCR in relevant clinical samples. In PBMCs from PwHD, repair associated DNA synthesis at *HTT* exon 1 showed a highly significant positive association with expanded CAG-repeat length and was readily detectable across the full range of disease-associated *HTT* CAG-repeat expansions. While the highest levels were observed in juvenile-onset HD, URSA was sensitive enough to detect a signal at CAG-repeat lengths below the clinical threshold of HD, including in an individual carrying an intermediate *HTT* allele in whom subtle HD-related phenotypes have been reported^52,65^. Notably, URSA also detected a signal in Stage 0–1 PwHD, demonstrating that activity of the DNA repair machinery is measurable before disease onset. Together, these observations indicate that URSA monitors the biological processes underlying SI rather than their cumulative outcomes. Unlike SE in blood, which accumulates slowly and requires years before significant repeat length changes become detectable, the rate of upstream repair activity driving instability can be quantified within days. Although the extent of somatic expansion and the effects of genetic modifiers can differ markedly across tissues^51^, key aspects of the SI mechanism including the influence of critical modifiers such as *MSH3*, *FAN1*, and *MLH3* are consistent between blood and brain^54,57^ and our findings further suggest that repair activity at the *HTT* DNA repair hotspots represents a shared, genetically driven biological process observable across both neurons and PBMCs. URSA scores in PBMCs correlated with the SER, which is influenced by both CAG-repeat length and age. Incorporating URSA into predictive models of somatic expansion improved their explanatory power suggesting that URSA captures individual variability in repair activity at the *HTT* that is not predicted by age and CAG-repeat length. Notably, this additional explanatory power relates to the individual-specific residual variance remaining after accounting for age and CAG repeat length, the same component that has been used to identify genetic modifiers of SI in blood^11^. Together, these results support the potential of peripheral measurement of repair at the m*HTT* exon 1 as a mechanistically informed biomarker for TE for SI modulating therapies with peripheral exposure. Further optimization will be required to adapt URSA for clinical application to use patient-derived samples under conditions that maintain drug exposure and TE.

A current design constraint of URSA is that repair activity is integrated over a minimum EdU incubation period of 24 hours. Since repair dynamics are likely considerably faster than this^66–68^, the assay cannot resolve whether one or multiple repair events occurred at a labeled locus within a given cell. In addition, the biochemical stringency required for EdU-DNA purification results in a conservative quantification, likely underestimating total repair activity. Despite this, repair frequency consistently exceeds expansion frequency across most analyzed PBMCs samples suggesting that most repair events are resolved without resulting in somatic expansion.

Genetic and experimental studies have implicated MMR in repeat instability in myotonic dystrophy, FECD3, and multiple spinocerebellar ataxias^51,69,70^. Our results provide direct evidence that frequent repair-associated DNA synthesis occurs at pathogenic repeats in human HD cells raising the possibility that other unstable expanded repeat loci may undergo similar repair dynamics, becoming DNA damage repair hotspots once their instability threshold is exceeded. The utility of URSA may extend further still, into oncology and other fields where modulating the DNA damage response is a therapeutic objective^71–77^.

In summary, URSA establishes a rapid, scalable and mechanistically grounded approach to quantify DNA repair activity at pathogenic repeat loci within days. By enabling direct measurement of repair engagement at the m*HTT* locus in both disease-relevant cell models and accessible patient-derived samples, this work provides a foundation for the development of minimally invasive TE biomarkers for SI–modulating therapeutics. Further optimization will be required before full implementation in clinical settings, and increased temporal resolution, sensitivity and throughput will be valuable to realize the complete potential of the assay in preclinical and translational applications.

**Supplementary Table 1:** Percentile ranking of the *HTT* exon 1 DNA repair signal relative to genome-wide DNA repair hotspots identified by URSA-seq. Percentile ranks were calculated by comparing the normalized URSA-seq signal at *HTT* exon 1 with all DRHs identified genome-wide within each cell line and EdU-labeling condition. Lower percentile values indicate a stronger repair signal and a higher ranking among genome-wide DRHs. Values are shown for RUES2-derived neurons, H9-derived neurons, and patient-derived iPSC neurons carrying the indicated *HTT* CAG repeat lengths.

**Supplementary Table 2. Baseline demographic and clinical characteristics of study participants.** Participants were stratified according to *HTT* CAG repeat length and clinical status. Values are presented as mean ± SD unless otherwise indicated. Expanded CAG repeat length is reported as the median (range). Individuals in HD-ISS stages 0-1 carry a pathogenic *HTT* expansion but had not developed clinical symptoms at the time of sampling. Age at onset is reported only for individuals in HD-ISS stages 2-3; data are not applicable for controls and participants classified as HD-ISS stages 0-1.

**Supplementary Table 3** Leave-one-out cross-validation (LOOCV) of regression models predicting SER. Predictive performance of the baseline regression model, including age at sampling, inherited CAG repeat length and their interaction (Age × CAG), was compared with the URSA-extended model, which additionally included URSA-mNR and its interaction terms (Age × CAG × URSA-mNR). Model performance was evaluated by leave-one-out cross-validation (LOOCV) using all 32 participants. Predictive accuracy is reported as the root mean squared error (RMSE) and mean absolute error (MAE) calculated from cross-validated predictions. Lower RMSE and MAE values indicate improved predictive performance. The LOOCV analysis demonstrates that inclusion of URSA-mNR improves prediction of SER beyond age and inherited CAG repeat length, supporting the robustness of the URSA-extended model.

## METHODS

### Research participants

Blood samples were obtained from participants recruited at the Movement Disorders Unit of Hospital de la Santa Creu i Sant Pau (Barcelona, Spain) (IIBSP-HUN-2017-105), as well as from additional control participants recruited at the Universitat de Barcelona (UB) and Vall d’Hebron Research Institute (VHIR). The study cohort comprised one intermediate allele carrier, 31 PwHD spanning a broad range of *HTT* CAG-repeat lengths up to juvenile-onset HD (>60 CAGs), representing Huntington’s Disease Integrated Staging System (HD-ISS) stages 0–3, and 25 unaffected non-mutation carrier controls of European ancestry (Supplementary Table 2). One individual carrying an intermediate allele (PBMCs35 CAGs) was included in the correlation analysis between URSA signal and CAG repeat length but was excluded from the control vs. HD diagnostic performance analysis due to the ambiguous clinical classification of intermediate alleles. The study protocol was approved by the Ethics Committee of Hospital de la Santa Creu iSant Pau, University of Barcelona and VHIR. Written informed consent was obtained from all participants prior to sample collection. All procedures were conducted in accordance with the ethical standards of the institutional and national research committees and with the 1964 Declaration of Helsinki and its subsequent amendments.

### Sample collection and PBMC isolation

Peripheral blood mononuclear cells were isolated from fresh whole blood collected from the study participants described above using standard Ficoll density gradient centrifugation in Cell Preparation Tubes (CPT; BD Biosciences, Cat. No. 362782). Samples were processed within 2 h of collection, and the PBMC fraction was recovered, washed in PBS containing 2% FBS, and resuspended in complete RPMI 1640 medium supplemented with FBS, L-glutamine, HEPES, and penicillin/streptomycin. Cells were counted and 2 × 10^6^ cells were seeded and incubated at 37°C with 5% CO₂. After 2 h, cells were treated with 20 μM EdU or DMSO as control. PBMCs were collected after 24 h by centrifugation.

### Cell lines

RUES2 ESC culture and neuronal differentiation. Human embryonic stem cells (ESCs) (RUES2 lines^49^: RUES2_20(22)CAG-cl.30 [CHDI-90001585], RUES2_56(20)CAG-cl.21.1 [CHDI-90001589] or RUES2_56(22)CAG-cl.23.1 [CHDI-90001590], RUES2_72(20)CAG-cl.12 [CHDI-90002877], and RUES2_118(22)CAG-cl.9 [CHDI-90004531]) were maintained in mTeSR1 (STEMCELL Technologies; Cat. No. 85850) on Geltrex-coated plates (Thermo Fisher Scientific; Cat. NoA1413302) and passaged every 4–5 days using Versene (Thermo Fisher Scientific; Cat. No15040066).

Cortical differentiation was performed by dual SMAD inhibition^78^ with minor modifications. ESCs were seeded in mTeSR1 supplemented with 10 µM Y-27632 and, upon confluency (DIV0), switched to N2-B27 medium containing 10 µM SB431542 and 100 nM LDN193189 for 8–9 days. N2-B27 comprised DMEM/F12 + GlutaMAX and Neurobasal (1:1) supplemented with N2 (0.5×), B27 (1×), insulin (2.5 µg/mL), MEM non-essential amino acids (1×), and 2-mercaptoethanol (100 µM). Neuroepithelial cells were dissociated with Dispase, replated onto poly-L-ornithine/laminin-coated flasks, and expanded in 20 ng/mL bFGF. At DIV15/16, neural progenitor cells were dissociated with Accutase and replated as single cells in N2-B27 medium supplemented with 10 ng/mL BDNF and GDNF. Cells were passaged again at DIV22/23 and plated at ∼200,000 cells/cm² on poly-D-lysine/laminin-coated plates. Differentiation was promoted with DAPT (γ-secretase inhibitor; Tocris, Cat. No. 2634) at 10 µM from DIV23/24–26/27, followed by 5 µM from DIV26/27–29/30. To inhibit proliferation, cultures were treated with 2 µM cytosine arabinoside (Ara-C; Tocris, Cat. No. 4520) for 48 h starting at DIV26/27. From DIV36/37, neurons were maintained in BrainPhys medium (STEMCELL Technologies; Cat. No. 05790) supplemented with BDNF, GDNF, ascorbic acid, cAMP, and laminin. Mature neurons were treated with 20 µM EdU (Sigma, Cat. No. #900584) for 3 or 7 days via half-medium changes; for 7-day treatments, EdU was replenished at DIV47/48 at half the final concentration. Cells were harvested at DIV50/51.

RUES2_118(22) CAG *MSH3* KO cells. The wild-type RUES2 cells contain 22 CAG repeats on one allele of the *HTT* gene and 20 on the other. To generate isogenic RUES2 cells that would be different only in the lengths of the CAG track of the *HTT* gene, a two-step CRISPR/Cas9-enabled gene editing approach was used. In the first step, a homology donor DNA plasmid (p*HTT*_Puro-TK) was constructed to insert a puromycin/thymidine kinase (PGK-Puro-TK) cassette to replace a 2248 bp *HTT* genomic region including part of its promoter (305 bp), the entire exon 1 (579 bp) and part of intron 1 (1364 bp). The two gRNAs used in this step were: 5’-gRNA: 5ʹ-GACGGGGACATTAGGCAGGCCGG-3ʹ, and 5’-gRNA: 5ʹ-GTTTAATTAGAAAATGATTCGG-3ʹ. The plasmid contained ∼1kb homology arms flanking the *HTT* genomic region to be replaced and a 23-bp gRNA target sequence derived from the zebrafish Tia1l gene on either side in opposite direction in-between the homology arms and PGK-Puro-TK cassette. In the second step, correctly edited RUES2 cells from step 1 containing a PGK-Puro-TK cassette on the *HTT*-CAG22 allele (or *HTT-*CAG20 allele) was used to re-insert the *HTT* genomic region with a 118 CAG repeat track but no other additional modifications back to the *HTT* gene, using the Tia1l-gRNA and a homology donor DNA plasmid containing the same homology arms as in p*HTT-*Puro-TK. Additionally, the *HTT* CAG118 donor DNA plasmid (p*HTT*_CAG118) reinstated the C/T SNP in the UTR region of exon 1. RUES2_118(20)CAG cells (CHDI-90004561), in which the original *HTT*-CAG22 allele was replaced with CAG118, were used to generate RUES2 CAG118; *MSH3*-/-cells. *MSH3* gene was deleted by CRISPR-induced non-homologous end-joining using a pair of upstream (intron 1: Intron1-gRNA1, GTAAAACTCTTCGACCACCAAGG or Intron1-gRNA2, ATTGAAACCAAGGCTGCTACCGG) and downstream (intron 8: Intron8-gRNA1, GGTACACTGGTACCCAACGGAGG or Intron8-gRNA2, CACAAAGGGTCACTAGGCTACGG) gRNAs that flanked a genomic region of approximately 40 kb that included *MSH3* exon 1 to exon 8.

H9 ESC culture and neuronal differentiation. Control (H9) and HD (CAG81) forebrain neuronal cells were generated and differentiated as previously described in ^50^ Briefly, the isogenic HD line carrying 81 CAG repeats (CAG81) was generated by TALEN-mediated editing of the H9 female human embryonic stem cell (hESC) line^79^. Cells were maintained under feeder-free conditions and differentiated into forebrain neurons through a neural progenitor cell stage followed by neuronal maturation, as previously reported. In this study, only the control H9 line and the HD CAG81 line were used, and genomic DNA from differentiated neuronal cells was provided for URSA assay analysis.

RPE-1^CAG129^ *MSH3* WT and *MSH3* KO cells. The human RPE-1 clonal cell model stably expressing *HTT*-exon 1 containing 129 CAG repeats (RPE-1^CAG129^) was previously described^33^ and *MSH3* knockout clones were generated by Balmus group (BioRxiv). RPE-1 cells were seeded at 1 × 10^6^ cells per 10-cm dish per condition and cultured in DMEM supplemented with 10% fetal bovine serum (FBS), 1% penicillin–streptomycin, 1% sodium pyruvate, and 1% L-glutamine at 37°C in a humidified atmosphere containing 5% CO₂. To induce G1-phase arrest, cells were treated with 250 nM Palbociclib 24 hours after seeding. After an additional 24 hours, a second dose of palbociclib was added together with 20 μM EdU or DMSO as vehicle control, and cells were incubated for a further 24 hours before collection for URSA assay analysis.

### Flow cytometry analysis

For EdU incorporation analysis, RPE-1 cells were seeded as described above and treated with EdU, with or without palbociclib, for 48 or 72 h. Cells were fixed with 4% paraformaldehyde and permeabilized before EdU detection using the Click-iT™ EdU Imaging Kit with Alexa Fluor™ 594 (Invitrogen, C10639) according to the manufacturer’s instructions. Samples were acquired on a Fusion II flow cytometer. Gating included exclusion of debris, selection of singlets and quantification of Alexa Fluor 594-positive cells. Fluorescence thresholds were established using DMSO-treated negative controls. Data were analysed using FlowJo (BD Biosciences).

PBMC proliferation was assessed by carboxyfluorescein succinimidyl ester (CFSE; Thermo Fisher Scientific) dilution. PBMCs were labelled with 1 μM CFSE for 5 min at 20 °C and cultured for 24 h. Cells were stained with Fixable Viability Dye eFluor™ 450 (eBioscience) and BV605-conjugated anti-human CD45 (clone HI30, BD Biosciences). Samples were acquired on a CytoFLEX flow cytometer (Beckman Coulter) and analysed using CytExpert v2.0 (Beckman Coulter). Cell proliferation was quantified from CFSE fluorescence dilution.

### Immunocytochemistry

Cells were fixed with 4 % PFA for 15 minutes at room temperature (RT), washed three times with PBS and blocked in 5 % Normal Goat Serum (NGS) + 0.3 % Triton-X-100 in PBS for 1 h at RT. After blocking, cells were incubated with primary antibodies overnight at 4°C, washed three times with PBS and incubated with secondary antibodies and DR for 1 hour at RT. After washes with PBS fluorescent images were acquired using an Opera Phenix high-content screening system (PerkinElmer) confocal microscope. To detect potential mitotic cells during the EdU treatment, the Click-iT EdU Imaging Kit (Thermo Fisher, Cat. No. C10337) was used after PFA fixation following the manufacturer’s instructions.

### Western blotting and capillary immunoassay

Cells were washed with PBS, PBS was removed and cells lysed in RIPA cell lysis buffer (Sigma, Cat. No. R0278-500ML) containing Protease Inhibitor (Roche Cat. No 04693159001)and Phosphatase Inhibitor (Roche, Cat. No. 04906837001) for 15 min on ice. Cells were scraped off the plate and centrifuged at 4°C for 10 min at 13,200 rcf. Protein concentration was measured using the Pierce™ BCA Protein Assay Kit (Thermo, Cat. No. 23227). A total of 10 µg protein per lane were separated on a 4-12% BisTris Bolt, 12-well SDS-PAGE gel and trasnferred onto a Immobilon-FL PVDF membrane (Millipore, Cat. No. IPFL00010).

#### Western blot

Blots were incubated overnight at 4°C with primary antibodies diluted in TBS-T with 5% Milk (anti-MSH3, abcam, Cat. No. ab275928, 1:300) or in Intercept Blocking Buffer (TBS) plus 0.1% Tween20 (anti-β-III Tubulin, R&D Systems/ Biotechne, Cat. No. MAB1195,1:20000). After washing, the blots were incubated for 1h at RT with secondary antibodies diluted 1:15000 in either TBS-T + 5% Milk (anti rabbit IRDye 800CW, LI-COR, Cat. No. 926-32211) or Intercept Blocking Buffer (TBS, LI-COR Bioscience) supplemented with 0.1% Tween-20 (anti mouse IRDye 800CW, LI-COR, Cat. No. 926-32210) depending on the primary antibody. Signals were detected on Odyssey® CLx Imager (LI-COR Biosicience) and quantified.

#### Capillary immunoassay

MSH3 protein expression was quantified using the automated Jess Simple Western system (Bio-Techne) with 12–230 kDa separation modules (Cat. No. SM-W004) according to the manufacturer’s instructions. Protein lysates were denatured at 95°C for 5 min before loading. Primary antibodies were anti-MSH3 (Abcam, ab275928; 1:300) and anti-β-actin (Cell Signaling Technology, Cat. No. 3700; 1:100). HRP-conjugated anti-rabbit (Bio-Techne, Cat. No. 043-426) and anti-mouse (Bio-Techne, Cat. No. 042-205) secondary antibodies were used for chemiluminescent detection. Data were acquired using the Jess instrument and analyzed with Compass software (Bio-Techne).

### Unscheduled DNA repair synthesis assay (URSA)

The URSA workflow builds on previously described enrichment and sequencing-based strategies for the detection of newly synthesized DNA, while incorporating optimized implementations for sequencing-based analysis (URSA-seq) and targeted locus analysis by digital PCR (URSA-dPCR) for quantitative analysis. Both approaches follow the core experimental principle established in Repair-seq^41^, based on EdU incorporation, genomic DNA fragmentation, click-mediated biotinylation, and streptavidin-mediated enrichment of newly synthesized DNA. For URSA-dPCR, biotinylated DNA fragments were directly quantified after enrichment by target-specific pre-amplification followed by digital PCR. For URSA-seq, fragmented DNA was subjected to end repair and adapter ligation prior to streptavidin enrichment, followed by library indexing, amplification, pooling, and high-throughput sequencing. The assay was implemented independently in two laboratories using closely related URSA workflows that shared the same overall experimental and analytical strategy, with protocol-specific adaptations in DNA extraction, fragmentation platform, click chemistry reagents, pre-amplification enzymes, library preparation, sequencing and dPCR instrumentation, and downstream bioinformatic processing.

### CAG-length measurement by capillary electrophoresis

Genomic DNA was extracted using the DNeasy Blood & Tissue Kit (QIAGEN #50969504). The expanded CAG tract was then amplified by PCR, using 150 ng of template DNA and the following primers: 6-FAM–labelled forward primer HD3F and reverse primer HD5 as reported in Collotta et al^33^. PCR products were analyzed on an ABI 3730xl Genetic Analyzer (Thermo Fisher Scientific), and fragment sizing was performed using GeneMapper software (Applied Biosystems/Thermo Fisher Scientific). Median CAG length, standard deviation, and instability index were calculated using the Romeo package as previously described.

### MiSeq sequencing of the *HTT* CAG repeat region

The *HTT* CAG repeat region was amplified and sequenced as described in Ciosi et al 2019^29^ (https://www.protocols.io/view/library-preparation-and-miseq-sequencing-for-the-g-rm7vzk6ervx1/v2). In brief, the repeat region was amplified from 20 ng genomic DNA using primers that amplify the repeat region and incorporate barcoded Illumina sequencing adapters. Sequencing was performed using the Illumina MiSeq platform at the University of Glasgow Shared Research Facility (SRF). Libraries were quantified using the KAPA library quantification kit and loaded at ∼1000 k clusters/mm^2^ supplemented with 5% PhiX DNA. Sequencing was performed using 400 bp forward and 200 bp reverse reads. Demultiplexed reads were aligned against custom reference sequences with variable CAG (1 to 200) and CCG (1 to 20) repeat lengths and common 5’-and 3’-flanking regions, and realigned where necessary to reference sequences with atypical allele structure. The inherited repeat length and allele structure for each allele were identified from the reference sequence that had the highest number of aligned reads. The alignment, genotype calling and calculation of SER10 (somatic expansion ratio 10) were performed using ScaleHD, an automated platform developed in-house (available on github at https://github.com/helloabunai/ScaleHD/). SER10 is the sum of reads at n+1 to n+10/number of reads at n, where n is the maximum number of reads aligned to any reference sequence. Aligned reads were visualised using Tablet to confirm correct alignment^80^.

## Statistical analyses

Statistical analyses were performed using GraphPad Prism (version 10.4.1) and R (version 4.5.2). Two-tailed Student’s *t*-tests and one-way analysis of variance (ANOVA) followed by Bonferroni or Tukey comparison test were performed in GraphPad Prism where appropriate. Linear mixed-effects models (LMMs), receiver operating characteristic (ROC) curve analyses, and multiple linear regression analyses were performed in R using the lme4, lmerTest, pROC, and stats packages, as appropriate. Details of statistical tests, sample sizes, and biological replicates are provided in the corresponding figure legends. Allele-specific URSA signal intensity across the CAG repeat length series was analyzed using linear mixed-effects models. To evaluate the association between URSA signal and pathogenic CAG repeat expansion, analyses were restricted to alleles within the pathogenic range (CAG ≥ 56). CAG repeat length was modeled as a fixed continuous effect, and experimental run (RUN) was included as a random intercept to account for batch-to-batch variability: URSAmt∼CAG+(1∣RUN). Degrees of freedom and *P* values were estimated using Satterthwaite’s approximation. ROC curve analyses were used to evaluate the ability of URSA to distinguish *HTT* expansion carriers from unaffected controls across progressively narrower expanded CAG repeat ranges. To determine whether URSA explained additional variation in SI beyond age and expanded CAG repeat length, ordinary least-squares multiple linear regression models were fitted using log-transformed somatic expansion ratio (logSER) as the dependent variable. A baseline model included age at sampling, expanded CAG repeat length and their interaction (logSER ∼ Age × CAG), whereas the extended model additionally included URSA-mNR (logSER ∼ Age × CAG × URSA-mNR). Because the baseline model is nested within the extended model, the models were compared using a nested ANOVA, with improvement in model fit assessed by the reduction in residual sum of squares (RSS) and changes in adjusted R².

## ACKNOWLEDGMENTS

We thank Gonzalo Olmedo-Saura for his valuable assistance with patient recruitment, clinical assessments, and coordination of sample collection. We thank Melanie Winkle and Susanne Luu-Heinicke for their expert technical support in generating the human neuron data. We thank Sara Tomaselli and Valentina Fodale for their assistance with cell culture models.

## DATA AVAILABILITY

The URSA-seq datasets generated during this study have been deposited in the European Nucleotide Archive (ENA) under accession numbers PRJEB115102 (H9 neurons, Brito laboratory) and PRJEB115090 (RPE1 CAG129, Balmus laboratory). Data generated by the CHDI Foundation are available upon reasonable request to the CHDI Foundation and subject to the Foundation’s data-sharing policies

## CONFLICT OF INTEREST DISCOLOSURE

G.B. is the founder and chief executive officer (part-time) of Function RX Ltd.

## Contributions

Conceptualization: P.B and V.B. Methodology: J.P.F., A.G., D.Y., C.C.E., M.B.G., A.V., D.R., P.P., V.B., and P.B. Investigation: J.P.F., S.A.C., J.F., K.v.B., K.H, D.Y., C.C.E., M.B.G., AL.B., K.P.L., N.F., H.L., G.M., L.G., and V.B. Formal analysis: J.P.F., A.G., S.A.C., A.B., N.F., M.M., A.V., A.I., D.M., and V.B. Data curation: S.A.C., A.L.B., A.I., M.M and V.B. Visualization: J.P.F., A.G., A.V., A.I., L.G., V.B., and P.B. Resources: J.P.P., M.C., F.C., C.V., R.Z.C., J.K., P.P., G.B., H.P.N., M.A.P., and D.M. Validation: L.G. Software: A.B. Project administration: A.G., J.P.P., V.B., and P.B. Supervision: J.P.P., B.P., T.F.V., G.B., H.P.N., M.A.P., D.M., V.B., and P.B. Writing – original draft: J.P.F., V.B., and P.B. Writing – review & editing: A.G., J.P.P., S.A.C., B.P., T.F.V., D.R., R.M.P., G.B., H.P.N., M.A.P., D.M., V.B., and P.B.

---

**Extended Data 1:**
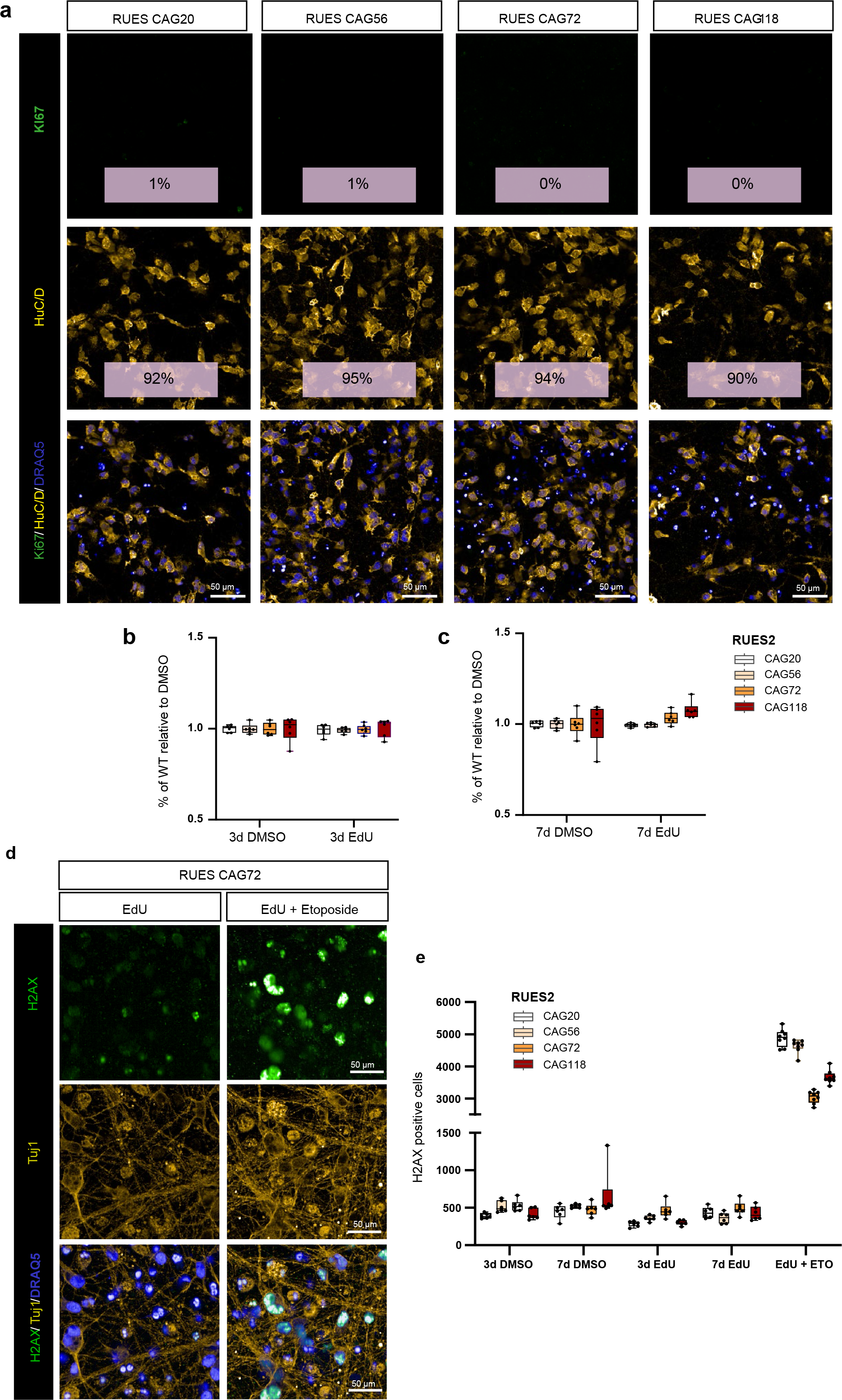
EdU labeling does not reflect proliferation and does not induce DNA damage in differentiated neurons. **(a)** Immunofluorescence of Ki67 (proliferation marker, green) and HuCD (neuronal marker, yellow) in neurons after 21 or 29 days of differentiation across RUES2 genotypes. Representative images show an absence of Ki67⁺ cells (∼0–1% positive cells), confirming post-mitotic cultures. Co-staining for HuC/D revealed a high proportion of neuronal cells (∼90–95%) across all experimental conditions, Scale bars 50 µm **(b, c)** Neuronal viability (percentage of live cells, normalized to DMSO control) across RUES2 genotypes after 3 **(b)** or 7 **(c)** days of treatment with DMSO or EdU (20 µM). Viability remained close to baseline (∼1.0) in all genotypes, indicating that prolonged EdU labelling is well tolerated. Box plots show the mean, with whiskers indicating minimum to maximum values. N=5 wells from one neuronal differentiation **(d)** Representative images of neurons stained for HuCD (yellow) and DRAQ5 (nuclear marker, blue) confirming neuronal identity and nuclear labeling. Scale bars 50 µm **(e)** Quantification of cell positives for the DNA double-strand break marker, γH2AX, across treatment conditions. Neurons treated with EdU for 3 or 7 days show γH2AX levels comparable to DMSO controls, indicating no increase in DNA damage upon EdU incorporation. Etoposide (ETO, positive control) induced a strong increase in γH2AX signal across all genotypes, confirming assay sensitivity. Box plots show the mean, with whiskers indicating minimum to maximum values. N=5 wells from one neuronal differentiation

**Extended Data 2:**
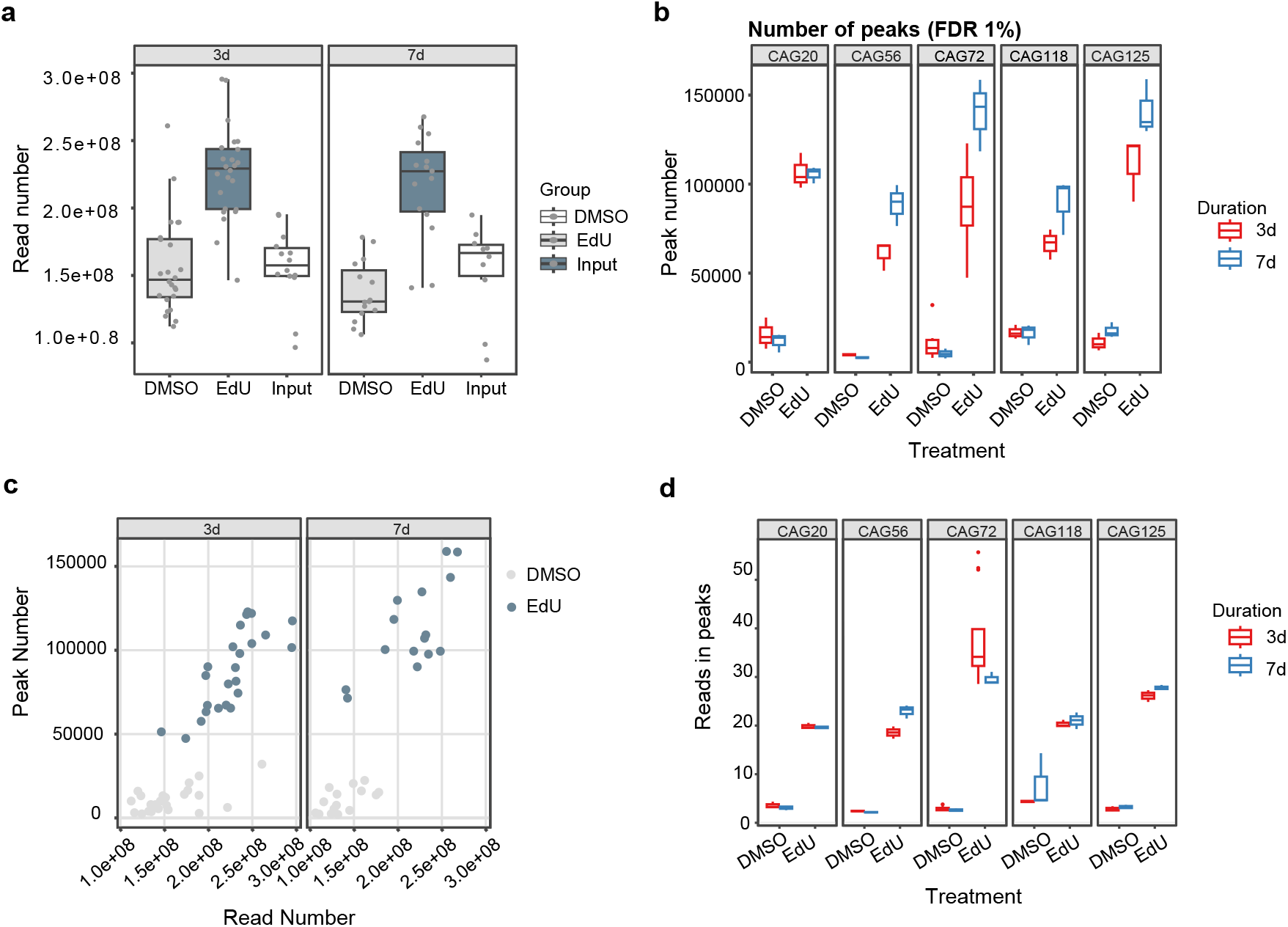
Sequencing quality control and peak-calling metrics. **(a)** Number of mapped paired-end reads across samples. Sequencing depth ranged from ∼80 to 300 million mapped reads per sample. **(b)** Number of peaks identified at FDR of 1% across RUES2 genotypes and labeling durations (3 and 7 days). Peak numbers vary modestly with sequencing depth. **(c)** Relationship between sequencing depth and peak number, indicating a moderate positive correlation. **(d)** Fraction of reads in peaks (FRiP) across samples.ranging from ∼15–35%, consistent with efficient enrichment.

**Extended Data 3:**
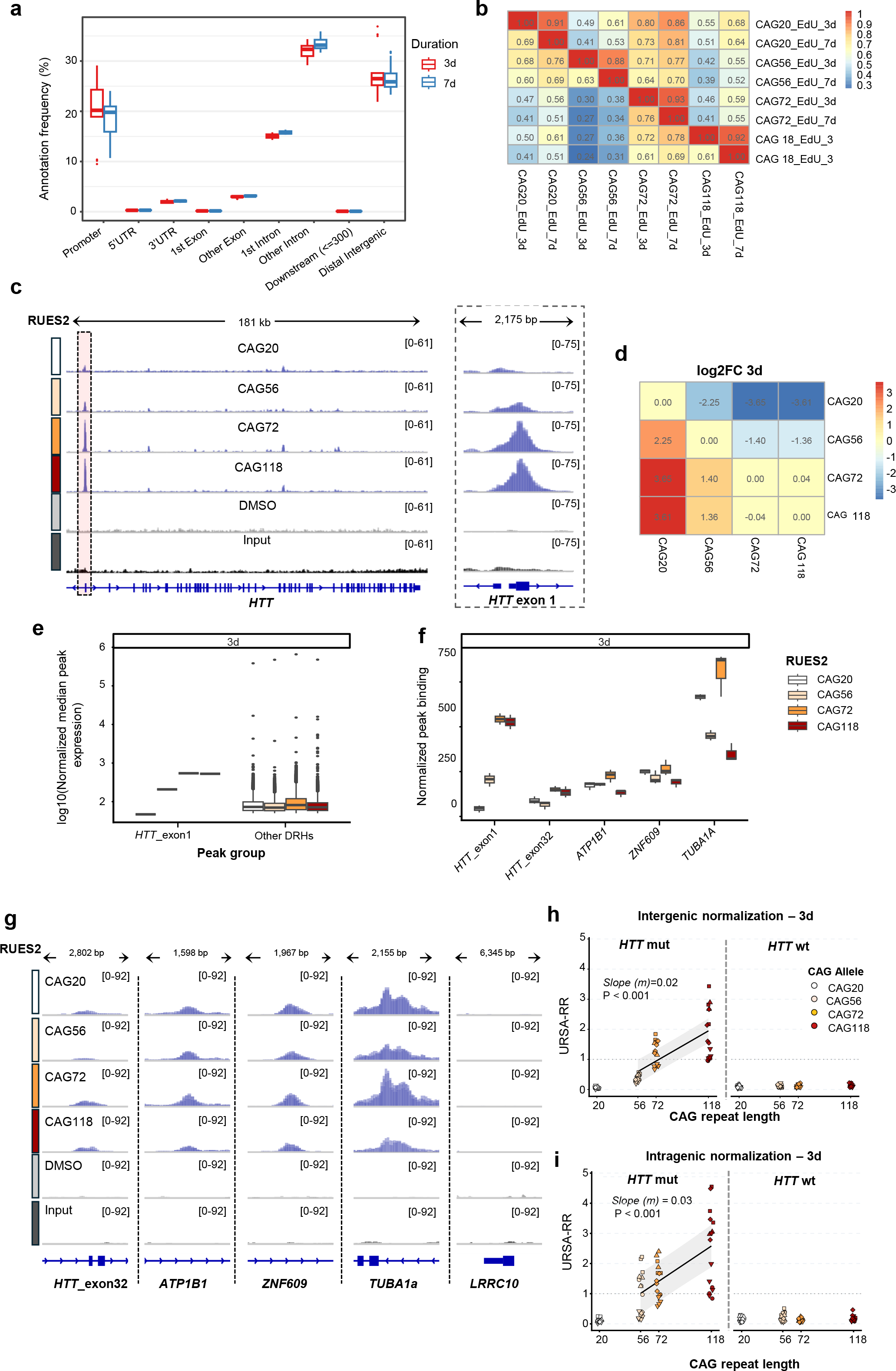
Genome-wide and locus-specific comparison of EdU-enriched regions following 3-and 7-day labeling in an allelic series of RUES2 derived neurons. **(a)** Distribution of URSA-seq peaks across genomic features (e.g., promoters, introns, intergenic regions) following 3-and 7-day EdU labeling, showing similar genomic annotations across labeling durations. **(b)** Proportion of shared peaks relative to the total peaks identified across samples and conditions,indicating reproducibility of URSA-seq peaks across labeling durations and genotypes. **(c)** IGV tracks of URSA-seq at the *HTT* locus following 3-day EdU treatment (20µM) across RUES2 genotypes (CAG20, CAG56, CAG72, CAG118) with input and DMSO controls. EdU-treated samples display clear enrichment across the *HTT* gene region, whereas input and DMSO controls show minimal background signal. N=3 wells from one neuronal differentiation **(d)** Heatmap of log₂ fold-change (log₂ FC) in EdU-enrichment at 3 days, normalized to input, across genotypes. **(e)** Comparison of normalized *HTT* exon 1 peak (*HTT*exon1) intensities against all other DRHs peaks (other_DRHs), by genotype after 3 days EdU incorporation. DRHs were defined as peaks called in at least two biological replicates per genotype, with ≥50 normalized counts, and absent in the DMSO control. Consistently with 7 days, *HTT* exon 1 signal is below median DRH intensity in WT neurons and among the highest in all HD genotypes **(f)** Normalized 3 day EdU incorporation signal at selected genomic loci, including *HTT* exon 1 and reference DRHs across RUES2 genotypes. **(g)** IGV tracks of URSA-seq at reference loci *ATP1B1*, *ZNF609*, *LRRC10* and *TUBA1A* after 3 day EdU treatment. Peak intensity is consistent across genotypes. N=3 wells from one neuronal differentiation **(h-i)** Allele-specific *HTT* exon 1 repair signal (URSA-RR) at the mutant (left) and wild-type (right) alleles across the CAG allelic series after 3 days of EdU treatment. URSA-RR was normalized to the geometric mean of *ATP1B1* and *ZNF609* (h) or to *HTT* exon 32 **(i)**. Each point represents one culture; symbol shape denotes the independent neuronal differentiation (n = 4) with 2–4 replicate cultures per differentiation. Solid lines in the mutant panels represent linear mixed-effects models with CAG repeat length as a fixed effect and neuronal differentiation as a random intercept (**f**: *m* = 0.02, P = 2.3 × 10⁻^12^; **g**: *m* = 0.03, P = 8.1 x 10^-10^). *m*: slope

**Extended Data 4:**
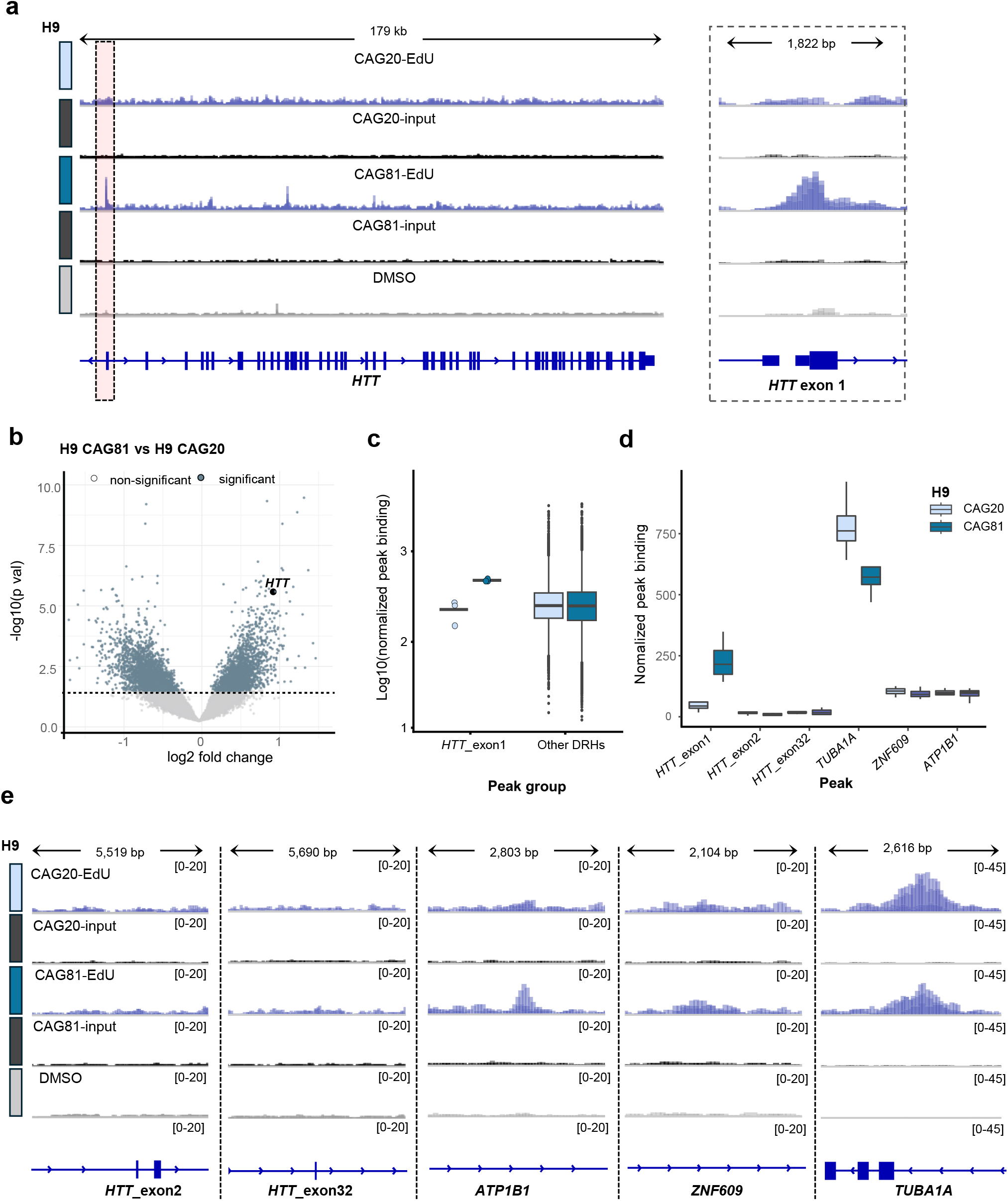
URSA-seq profiling reveals substantial EdU incorporation at the HTT exon 1 locus in H9 derived neurons. (a) IGV tracks of URSA-seq at HTT locus following 7-day EdU treatment (20 µM) in WT H9 CAG20 and HD H9 CAG81 neurons with input and DMSO controls. EdU treated H9 CAG81 samples display clear enrichment at the exon 1 of HTT, whereas WT, input, and DMSO controls show minimal background signal. The red-boxed region marks HTT exon 1, shown in the zoomed panel to the right. N=3-4 wells from 3 independent neuronal differentiations (b) Volcano plot showing differential enrichment analysis of URSA-seq peaks for pairwise comparisons of HD vs WT H9 cell lines (CAG81 vs CAG20). Each point represents a genomic peak. Peaks meeting significance thresholds (FDR < 0.01 and log₂ fold-change > 4) are shown in green (N = 3-4/genotype). The HTT exon 1 DRH identified in (a) is annotated. (c) Log₁₀-normalized median EdU enrichment at HTT exon 1 versus all other DRHs, by genotype. DRHs were defined as peaks present in ≥2 biological replicates per genotype, with ≥50 normalized counts, and absent in DMSO controls. HTT exon 1 signal falls below median DRH intensity in WT cells but is among the highest in CAG81 neurons. (d) Normalized EdU signal at HTT exon 1 relative to other DRHs loci across genotypes. Points represent biological replicates; error bars denote mean ± s.e.m, (n=3-4 per genotype) (e) IGV tracks of reference loci TUBA1A and ZNF609 in H9 cells. TUBA1A shows a clear peak consistent with RUES2 neurons, ZNF609 shows a weaker but stable peak. N=3-4 wells from 3 independent neuronal differentiations.

**Extended Data 5:**
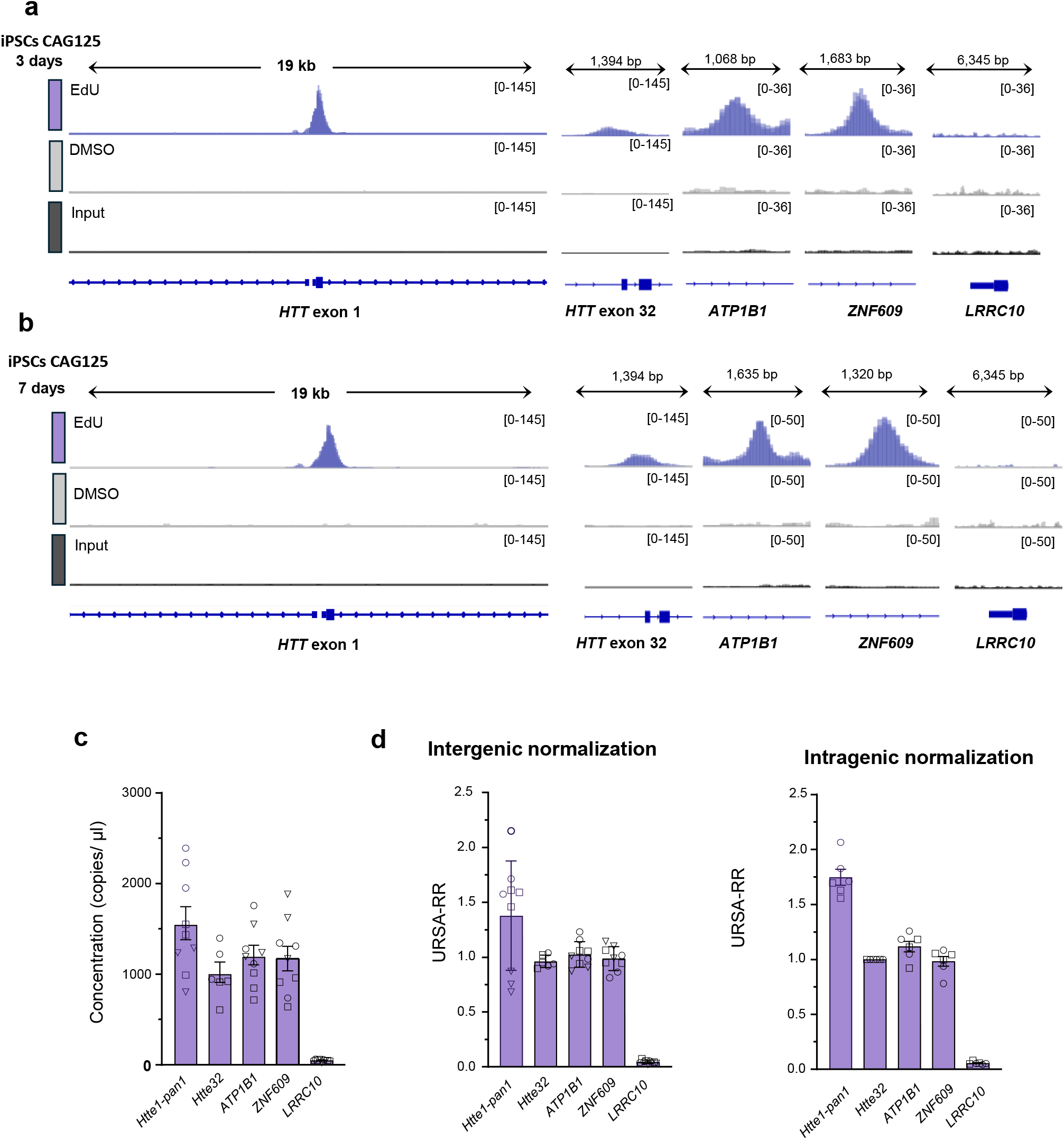
URSA-seq reveals substantial EdU incorporation at the HTT exon 1 locus in iPSCs CAG125 derived neurons. (a, b) IGV tracks at the HTT locus in HD CAG125 (CAG125_EdU) compared to input and DMSO control (CAG125_DMSO). A focal EdU enrichment peak is present at HTT exon 1 after 3 (a) and 7 (b) days of EdU treatment. N=3 experimental runs, i.e., number of independent neuronal differentiations (c) URSA-ddPCR raw signal (copies/µL) at HTT exon1 using non-allele specific primers (HTTe1-pan1), HTT exon 32, and ATP1B1, ZNF609 and LRRC10 after 7 days of EdU incorporation. N=3-4 wells from 3 independent neuronal differentiations (different shapes). (d) Normalized URSA-dPCR (URSA-RR) signal at the HTT exon 1 DRH using intergenic normalization to geometric mean of ATP1B1 and ZNF609 (left panel) and intragenic normalization to HTT exon 32 (right panel). Bars represent mean ± s.e.m. across biological replicates, N=3-4 wells from 2-3 independent neuronal differentiations (different shapes)

**Extended Data 6:**
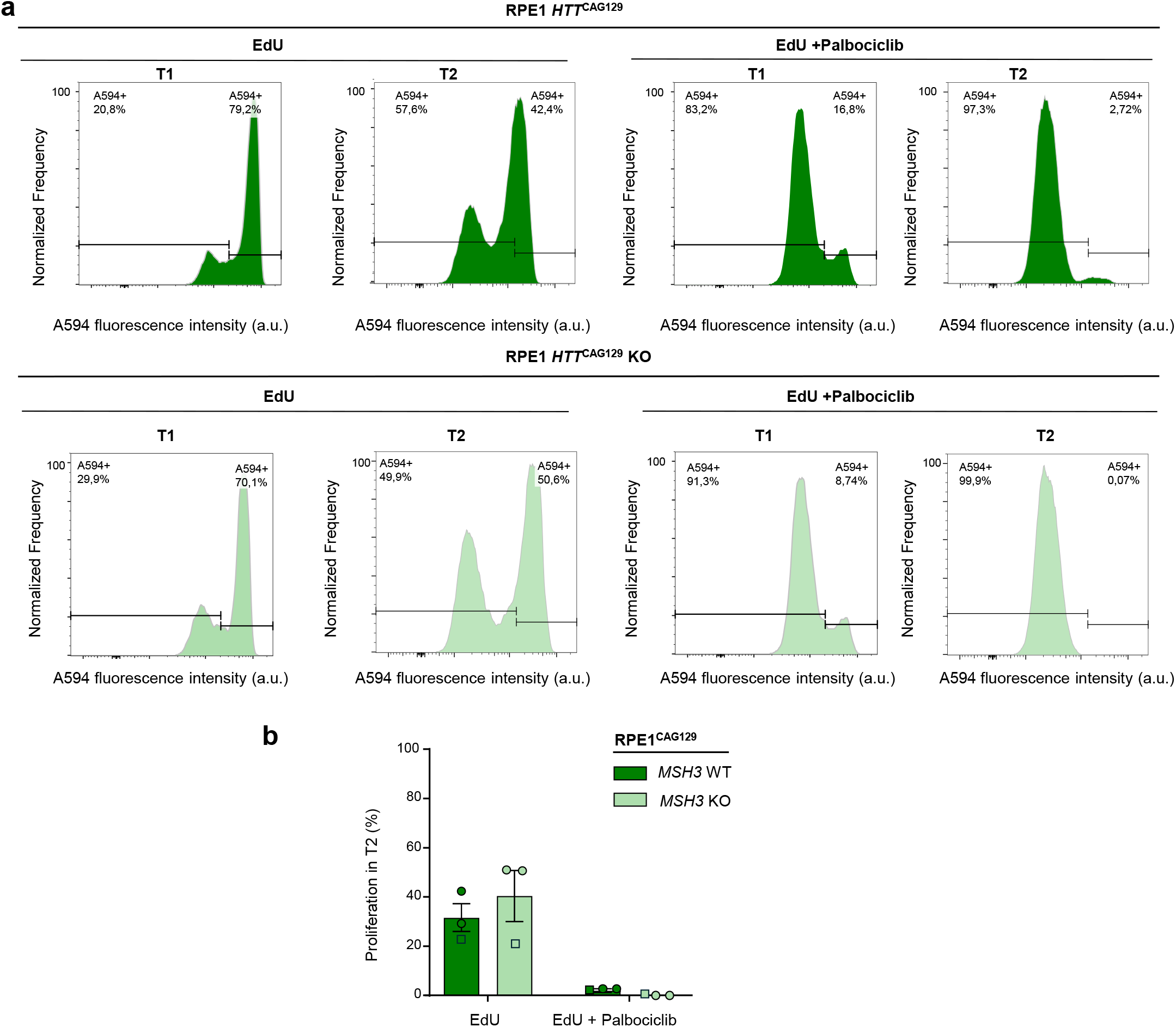
Validation of G1-phase arrest and absence of proliferation in RPE-1 HTT cells. **(a)** Representative flow cytometry plots of EdU incorporation (Click-iT, Alexa Fluor 594) in RPE-1 *HTT*CAG129 and RPE-1 *HTT*CAG129 MSH3-KO cells treated with EdU alone or with EdU plus palbociclib. EdU-treated cells showed a substantial population of Alexa Fluor 594-positive cells, indicating active DNA replication, whereas combined EdU and palbociclib treatment markedly reduced EdU incorporation, consistent with G1-phase arrest. **(b)** Quantification of Alexa Fluor 594-positive cells confirmed a strong reduction in EdU-positive cells following palbociclib treatment in both WT and MSH3-KO cells, supporting effective inhibition of S-phase entry prior to URSA analysis. Bars represent the mean ± s.e.m. of two independent experiments. Individual points represent wells and different shapes represent independent runs. (N = 2 independent experiments; 3 wells per experiment per genotype).

**Extended Data 7:**
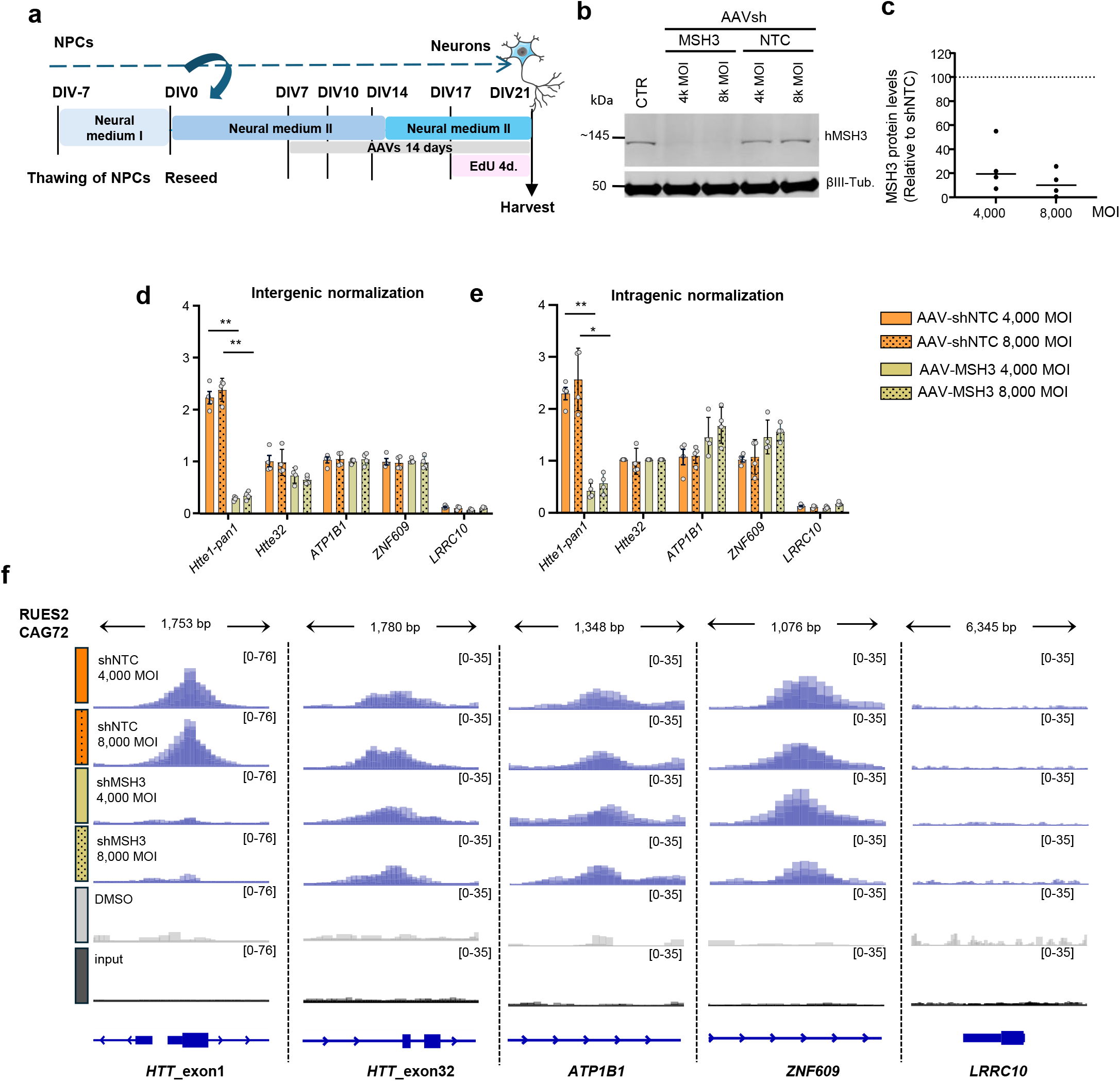
URSA detects MSH3 modulation by AAV-delivered shRNA. **(a)** Schematic of neuronal differentiation, AAV-shRNA transduction timing, and EdU labeling. RUES2-72CAG neurons were transduced with shRNA at the indicated time point and exposed to EdU for 4 days prior to collection. (**b)** Representative Western blot confirming MSH3 knockdown by shRNA in CAG72 neurons. **(c)** Quantification showing robust reduction in MSH3 protein levels upon shRNA treatment. **(d, e)** Intergenic (**d**) and intragenic (**e**) normalized URSA-dPCR signal (URSA-RR) at *HTT* exon 1 following transduction with non-targeting (AAV-shNTC) or *MSH3* targeting (AAV-MSH3) shRNA at 4000 MOI or 8000 MOI, scaled to the non targeting condition. Only 13% to 22% of the URSA-dPCR signal at the HTT exon1 locus is left after MSH3 knock-down. Dots represent biological replicates; error bars denote mean ± s.e.m. N=4 wells from one neuronal differentiation). * P < 0.05; **P < 0.01, one-way ANOVA **(f)** IGV tracks of URSA-seq at *HTT* exon 1 in CAG72 RUES2 neurons transduced with either MSH3 shRNA (AAV-shMSH3) or non-targeting shRNA (AAV-shNTC) compared to input (input) and DMSO (DMSO). N=3 wells from one neuronal differentiation

**Extended Data 8:**
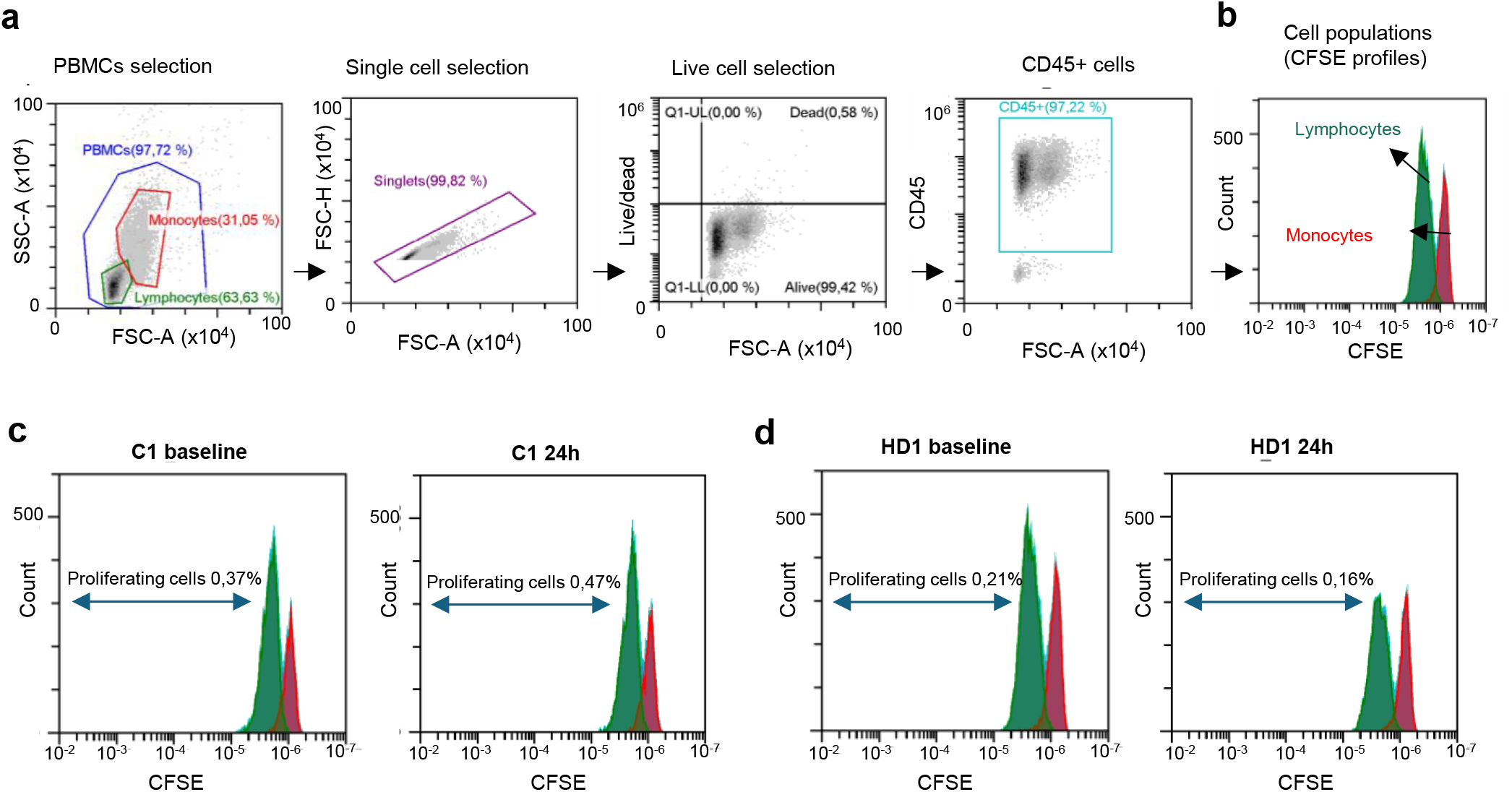
CFSE-based analysis of PBMC proliferation. **(a)** Representative scatter plots showing gating strategy: PBMCs (lymphocytes and monocytes) were defined by side-scatter area (SSC-A) and forward scatter area (FSC-A). FSC-Height vs FSC-Area were used to exclude cell aggregates from the analysis. Dead cells were excluded with a fixable viability stain. Total leukocytes were selected by gating on CD45+ cells. **(b)** Representative scheme illustrating identification of proliferating CD45⁺ cells based on CFSE signal decay. Proliferation of lymphocytes and monocytes was assessed separately. **(c, d)** Representative histograms from **(c)** a control individual and **(d)** an HD patient at baseline (day 0) and after 24 h show minimal CFSE dilution, indicating negligible proliferation over this time frame.

**Extended Data 9:**
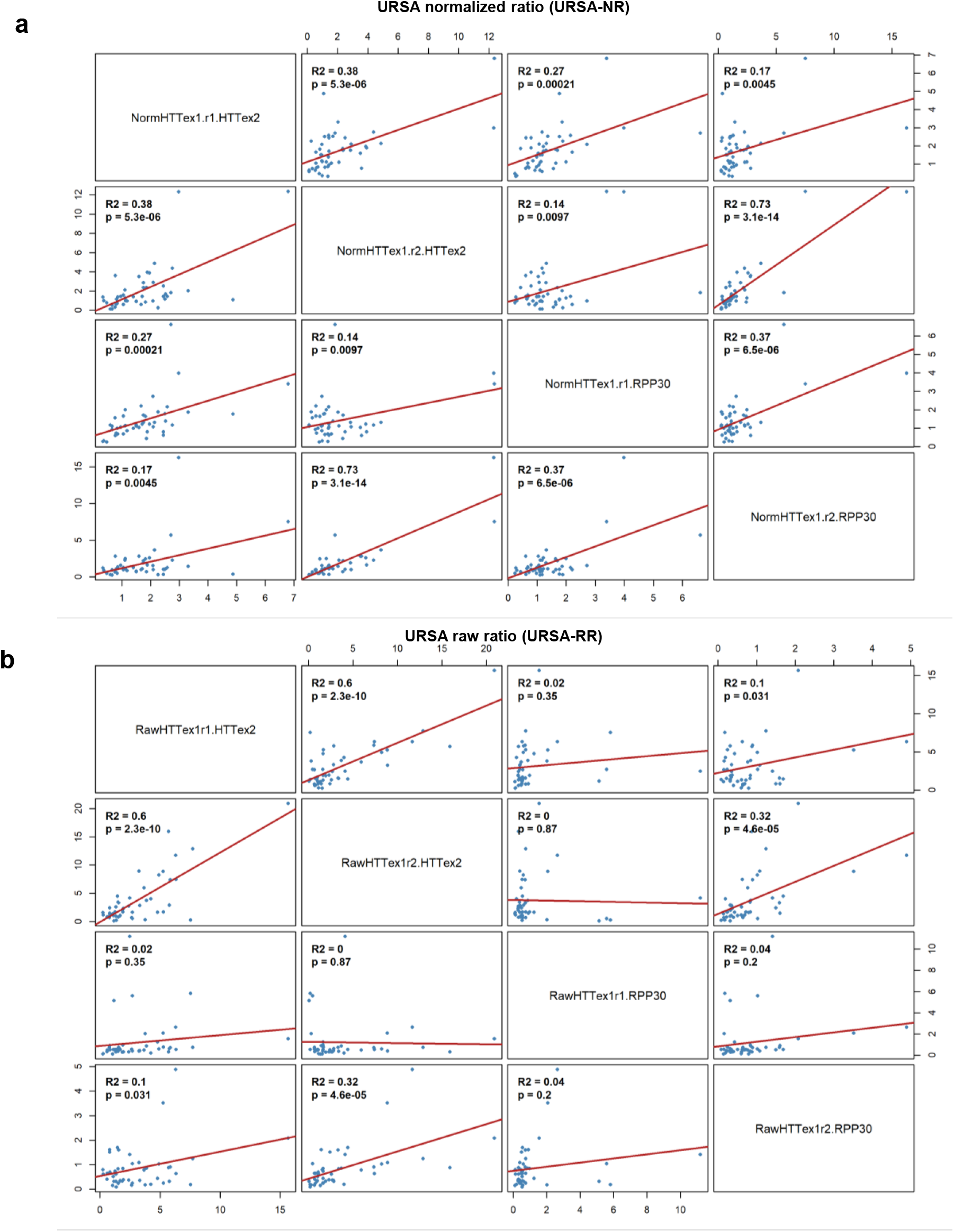
Comparison of normalization strategies for URSA-dPCR measurements. (a) Pairwise correlations between URSA normalized ratios (URSA-NR) obtained using two independent HTT exon 1 primer sets and normalized to either an intragenic (HTT exon 2) or an intergenic (RPP30) reference, following batch-relative scaling to the mean of control samples within each experimental batch. (b) Pairwise correlations between corresponding raw ratios (URSA-RR) without batch scaling. Analyses include all controls (N=21 including one non-HD individual with 35 CAG repeat) and HD samples (N=31). Batch-scaled values show higher concordance across normalization strategies than raw ratios. Each dot represents an individual sample; red lines indicate linear regression fits, with coefficients of determination (R²) and p-values shown in each panel.

