## Supplementary Tables for "Unscheduled DNA synthesis reveals a DNA repair hotspot and biomarker of somatic instability at the expanded-CAG repeat tract in the huntingtin gene"

**Supplementary Table 1:** Percentile ranking of the *HTT* exon 1 DNA repair signal relative to genome-wide DNA repair hotspots identified by URSA-seq.

| | RUES2 | | | | H9 | | iPCS pwHD |
|---|---|---|---|---|---|---|---|
| | CAG20 | CAG56 | CAG72 | CAG118 | CAG20 | CAG81 | CAG125 |
| 3 days | - | 1 | 0.12 | 0.05 | | | 0.01 |
| 7 days | 91,5 | 3.3 | 0.05 | 0.03 | 85 | 0.3 | 0.01 |

**Supplementary Table 2.** Baseline demographic and clinical characteristics of study participants

| Group | Control <27 CAG | Intemediate allele carrier | *HTT* expanded allele carriers across HD-ISS stage*s* | |
|---|---|---|---|---|
| | | | **0 - 1** | **2 - 3** |
| **n** | 24 | 1 | 6 | 35 |
| **Age at sampling,years** | 42.6 ± 18.6 | 70 | 45.8 ± 9.5 | 50 ± 16.2 |
| **Sex (Male/Female)** | 14 / 10 | 0/1 | 3 / 3 | 11 / 14 |
| **CAG repeats (median)** | - | 35 | 41.5 (39-43) | 45 (40-73) |
| **Age at onset, years** | - | 27 | N/A | 40.4 ± 14.9 |

**Supplementary Table 3.** Leave-one-out cross-validation (LOOCV) of regression models predicting somatic expansion ratio (SER).

| Metric | Baseline model (Age × CAG) | URSA-extended model (Age × CAG × URSA-mNR) |
|---|---|---|
| Sample size (n) | 32 | 32 |
| LOOCV RMSE | 0.3693 | 0.1515 |
| LOOCV MAE | 0.2279 | 0.1167 |
